# Cytosolic condensates enriched in polyserine repeats are preferred sites of tau fiber propagation

**DOI:** 10.1101/2022.09.22.509111

**Authors:** Evan Lester, Meaghan Van Alstyne, Kathleen L. McCann, Spoorthy Reddy, Li Yi Cheng, Jeff Kuo, James Pratt, Roy Parker

**Author notes:** Contributed equally (Evan Lester, Meaghan Van Alstyne).

## Abstract

Tau aggregates are a hallmark of multiple neurodegenerative diseases and can contain RNAs and RNA binding proteins, including SRRM2 and PNN. How these resident nuclear proteins mislocalize and their influence on the prion-like propagation of tau fibers remains unknown. We demonstrate that polyserine repeats in SRRM2 and PNN are necessary and sufficient for recruitment to tau aggregates. Moreover, we demonstrate tau fibers preferentially grow in association with endogenous cytoplasmic assemblies – mitotic interchromatin granules and cytoplasmic speckles – which contain SRRM2 and PNN. Polyserine undergoes self-assembly *in vitro* and in cells, where polyserine-assemblies are sites of tau fiber propagation. Modulating the levels of polyserine containing proteins results in a corresponding change in tau aggregation. These findings define a specific protein motif, and cellular condensates, that promote tau fiber propagation. As cytoplasmic speckles form in iPSC neurons under inflammatory or hyperosmolar stress, they may promote tau fiber propagation in various neurodegenerative diseases.

## Introduction

Tau inclusions are a pathological hallmark defining over 20 neurodegenerative diseases including Alzheimer’s disease (AD), corticobasal degeneration (CBD), and hereditary frontotemporal dementia with parkinsonism-17 (FTDP-17), collectively classified as tauopathies (Wang and Mandelkow, 2016). While loss-of-function mechanisms can contribute to disease, tau knockout mice have no overt neurodegenerative phenotype and show mild deficits at advanced ages (Ke et al., 2012). Instead, several observations argue that the formation and spread of tau oligomers or aggregates can be causative of neurodegenerative disease. For example, mutations in familial FTDP-17 promote tau aggregation (Spillantini et al., 2000), induction of tau aggregates leads to toxicity in cells (Sanders et al., 2014), and targeted reduction of tau and inhibition of tau aggregation reverses cognitive impairments in tauopathy mouse models (Anglada-Huguet et al., 2021; DeVos et al., 2017, 2018). The aggregation of tau has been proposed to propagate through prion-like mechanisms originating via misfolding of an initial seed and progressing into larger, fibrillar inclusions (Goedert and Spillantini, 2017). Nonetheless, the mechanistic basis for tau-mediated aggregation and neurodegeneration remains incompletely understood.

A key area of interest in the formation of tau fibers and aggregates are the molecules and subcellular locations that modulate tau fiber formation and propagation. Previous work has shown that tau binds microtubules and pathogenic mutations or post-translational modifications reduce tau’s interaction with microtubules leading to increased tau fiber formation (Iqbal et al., 2016). Yet, *in vitro* tau fibrillization typically requires a polyanionic co-factor such as RNA or heparin (Dinkel et al., 2015; Friedhoff et al., 1998; Kampers et al., 1996). This *in vitro* requirement suggests that tau fiber formation in cells will require cofactors, which remain unknown. Interestingly, tau can form condensates *in vitro* with RNA (Zhang et al., 2017) suggesting an RNA containing condensate within cells might contribute to tau fiber formation and/or propagation, although direct evidence for a specific cellular condensate promoting tau aggregation is lacking.

We recently showed that nuclear tau aggregates, which can be observed in cell lines, mouse models, or human post-mortem samples, are observed in association with nuclear speckles (Lester et al., 2021). Nuclear speckles are a condensate made up of nascent transcripts and components of the transcription or RNA processing machinery (Spector and Lamond, 2011). Moreover, we observed mislocalization of some nuclear speckle proteins to cytosolic tau aggregates (Lester et al., 2021). Two of the most prominently mislocalized proteins are pinin (PNN) and serine/arginine repetitive matrix protein 2 (SRRM2). The mislocalization of SRRM2 to cytoplasmic tau aggregates is relevant to disease since this occurs in tau mouse models of disease and tauopathy patient brains (Lester et al., 2021; McMillan et al., 2021). Congruent with these findings, SRRM2 is an RNA binding protein involved in RNA splicing and RNA sequencing shows tau aggregation induces RNA splicing changes in several model systems (Apicco et al., 2019; Lester et al., 2021), as well as in patients with AD (Hsieh et al., 2019; Raj et al., 2018). These features liken tauopathies to other neurodegenerative diseases in which dysfunction and mislocalization of RNA binding proteins (RBPs) leads to dysregulated RNA processing and gene expression (Conlon and Manley, 2017). However, how the interaction between nuclear speckle components and tau aggregates contributes to aggregate formation and/or toxicity remains unknown.

Here, we investigated the mechanisms mediating mislocalization of nuclear speckle proteins to tau inclusions and their involvement in tau aggregation. We identified polyserine stretches in SRRM2 and PNN that contribute to, and are sufficient for, recruitment to tau aggregates. We also identified cytoplasmic condensates containing SRRM2, referred to as mitotic interchromatin granules (MIGs) and cytoplasmic speckles (CSs), that serve as preferential sites of tau aggregate growth. Importantly, decreasing the polyserine containing protein PNN, or overexpressing polyserine domains, correspondingly decreases or increases tau aggregation. These findings delineate homopolymeric serine stretches as a mediator of protein recruitment to tau aggregates as well as a defining feature of assemblies that specify a preferred subcellular location for tau aggregation.

## RESULTS

### The C-terminal regions of SRRM2 and PNN mediate association with tau aggregates

Our previous work suggested the long C-terminal disordered region of SRRM2 was required for its mislocalization to cytoplasmic tau aggregates (Lester et al., 2021). To determine if there is a specific motif that dictates SRRM2-tau interactions, we used HEK293 tau biosensor cells to generate a series of cell lines with truncated SRRM2 by inserting Halo tags into chromosomal copies of the SRRM2 gene using the CRISPaint system (Ilik et al., 2020; Schmid-Burgk et al., 2016). The HEK293 tau biosensor cells express both cyan fluorescent protein (CFP) and yellow fluorescent protein (YFP) tagged forms of the tau K18 fragment with the P301S mutation and form bright FRET+ (Förster resonance energy transfer) tau aggregates upon lipofection of exogenous tau seeds isolated from the brains of Tg2541 (P301S) tauopathy mice (Holmes et al., 2014). All endogenous Halo-tagged SRRM2 truncations were detectable at the appropriate size (Figure 1A, S1A). The multiple bands observed in truncations 5-7 could represent unannotated splice isoforms of SRRM2 or post-translational modifications, which are revealed because of the smaller size of these truncation proteins (Figure S1A).

**Figure 1:**
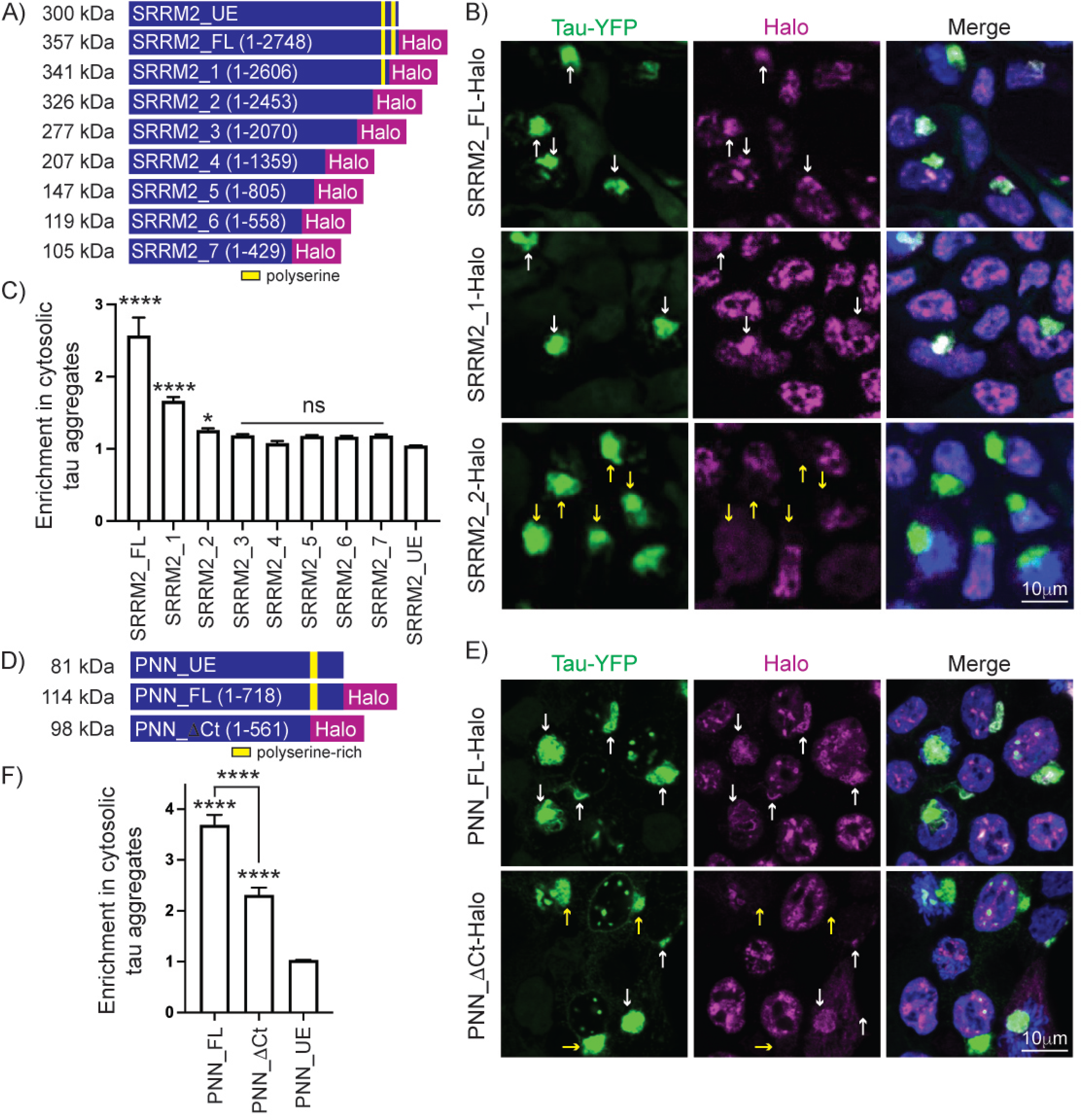
SRRM2 and PNN C-termini mediate enrichment in tau aggregates. **(A)** Schematic of SRRM2 truncations generated through CRISPaint in HEK293 tau biosensor cells where Halo tags were introduced into endogenous SRRM2 loci followed by a polyadenylation signal to create tagged and truncated proteins. Yellow regions denote two polyserine stretches in the SRRM2 C-terminus **(B)** Immunofluorescence of tau-YFP (green), Halo (magenta), and DAPI (blue) in SRRM2_FL-Halo, SRRM2_1-Halo, and SRRM2_2-Halo lines showing enrichment of SRRM2 (white arrows) or lack of enrichment (yellow arrows) in tau aggregates seeded via lipofection of clarified brain homogenate from tau transgenic mice (Tg2541). Images of all truncations can be found in Fig S2A. **(C)** Quantification of the ratio of Halo mean intensity within tau aggregates relative to Halo mean intensity in the cytoplasm for SRRM2 truncations. n=75 images quantified from three biological replicates per group. Data represent mean and 95% CI. Statistics performed with one-way ANOVA with comparison to SRRM2_UE. (*) P < 0.05 (****) P < 0.0001. **(D)** Schematic of PNN truncation made through CRISPaint introducing a Halo tag at the endogenous PNN loci. **(E)** Representative images of Halo labeling (magenta) and tau-YFP (green) in full-length Halo tagged PNN (PNN_FL-Halo) and C-terminal truncated PNN (PNN_DCt-Halo) cell lines showing enrichment of PNN (white arrows) or no enrichment (yellow arrow) in tau aggregates. **(F)** Quantification of unedited (PNN_UE), PNN_FL-Halo, and PNN_DCt-Halo enrichment in cytoplasmic tau aggregates as in (C). Data represent mean and 95% CI. n>788 cells from three biological replicates per group. Statistics performed with one-way ANOVA. (****) P < 0.0001.

This analysis demonstrated that the last 294 amino acids in the C-terminal region of SRRM2 are necessary for recruitment to tau aggregates. Specifically, we observed deletion of the last 146 amino acids (SRRM2_1, a.a. 2606-2752) significantly reduced enrichment in tau aggregates (from a mean enrichment of 2.57 for full length SRRM2 to 1.67) while a larger 294 amino acid deletion (SRRM2_2, a.a. 2458-2752) led to a near complete loss of enrichment (mean enrichment of 1.26) (Figure 1C). All larger deletions failed to co-localize with tau aggregates (Figure 1C, S2A). Furthermore, SRRM2 truncations 6 and 7 display a diffuse nuclear localization indicating amino acids 558-805 are required for normal recruitment of SRRM2 to nuclear speckles (Figure S2A).

The amino acid sequence of SRRM2 has two homopolymeric serine stretches in the C-terminal 294 amino acids, one that is 42 serines long and a second that is 30 serines long (Figure 1A, yellow boxes). The SRRM2_1 truncation removes the 42-polyserine stretch while the SRRM2_2 truncation removes both the 42 and 30-polyserine stretches. These results suggested two regions within the C-terminus of SRRM2, each containing a polyserine domain, are required for enrichment of SRRM2 in tau aggregates (see below).

The nuclear speckle protein PNN also mislocalizes to cytoplasmic tau aggregates (Figure S1C) (Lester et al., 2021). The C-terminus of PNN contains a region of 53 amino acids of which 44 (83%) are serine residues. To determine whether this serine-rich region of PNN mediates associations with tau aggregates, we utilized CRISPaint to generate cell lines expressing Halo-tagged full-length PNN (PNN_FL) or a truncated (PNN_ΔCt) form with deletion of the terminal 157 amino acids including the serine-rich region (Figure 1D, S1B). We observed the C-terminal truncation of PNN resulted in a significant reduction in recruitment to tau aggregates compared to the full-length protein (Figure 1E, F), providing additional evidence that serine-rich regions mediate associations with tau aggregates.

Taken together, these findings identify polyserine or serine-rich domains as elements resulting in mislocalization and recruitment of two nuclear speckle proteins to tau aggregates. Interestingly, there are fewer than 20 human proteins which contain pure stretches of serine longer than 20 amino acids and several of the top serine-repeat containing proteins are involved in RNA homeostasis (Supplemental Table 1). In addition to SRRM2 and PNN, we tested SETD1A – a histone methyltransferase implicated in neurodevelopmental disorders that contains 24 consecutive serines – and observed enrichment in tau aggregates (Figure S1D).

### Serine-rich protein domains, or polyserine alone, are sufficient for association with tau aggregates

To determine if the polyserine containing regions of SRRM2 and PNN are sufficient for recruitment to tau aggregates, we exogenously expressed Halo-tagged C-terminal SRRM2 fragments containing both polyserine regions (Frag_2: a.a 2458-2752), one polyserine region (Frag_1: a.a. 2606-2752), or no polyserine regions (Frag_0: a.a. 2651-2752) (Figure 2A). We also expressed amino acids 561-637 from PNN fused to Halo, where 50 out of 77 amino acids are serine residues (Figure 2D).

**Figure 2:**
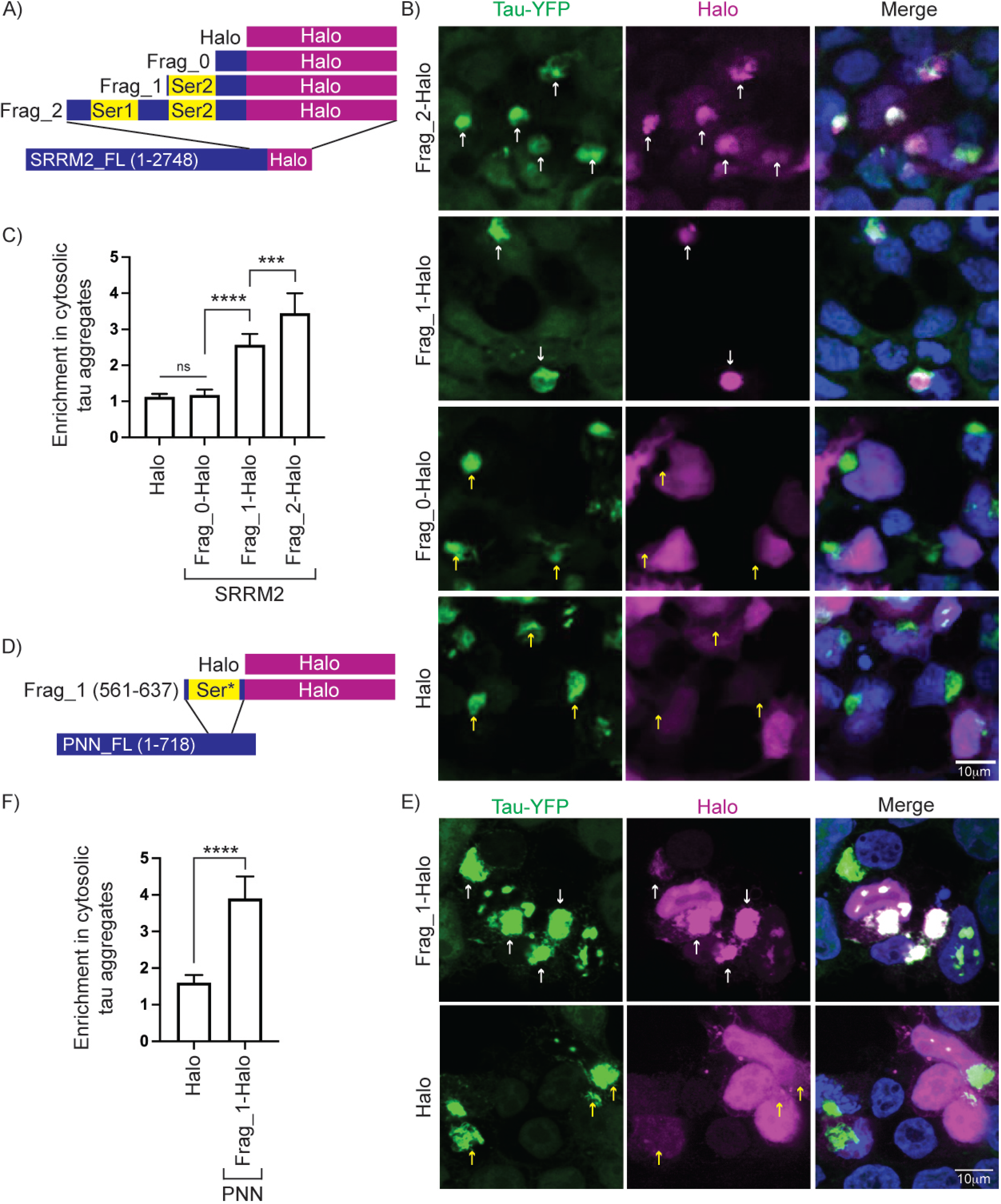
SRRM2 and PNN C-termini are sufficient for localization to tau aggregates. **(A)** Schematic of Halo tagged SRRM2 C-terminal fragment constructs and control. **(B)** Immunofluorescence of tau-YFP (green), Halo (magenta), and DAPI (blue) in HEK293 biosensor cells transfected with constructs in (A) and seeded with clarified brain homogenate showing colocalization with tau aggregates (white arrows) and lack of colocalization (yellow arrows). **(C)** Quantification of the ratio of mean intensity of Halo within cytoplasmic tau aggregates relative to Halo signal in the surrounding cytoplasm for SRRM2 C-terminal fragments. n=25 tau aggregates per group. Data represent mean and 95% CI. **(D)** Schematic of construct encoding a Halo tagged serine-rich fragment of PNN. **(E)** Immunofluorescence of tau-YFP (green), Halo (magenta), and DAPI (blue) in HEK293 biosensor cells transfected with constructs in (D) and seeded with clarified brain homogenate showing colocalization with tau aggregates (white arrows) and lack of colocalization (yellow arrows). **(F)** Quantification of the ratio of mean intensity of Halo within cytoplasmic tau aggregates relative to Halo signal in the remaining cytoplasm for PNN C-terminal fragment. n>140 cells from three biological replicates per group. Data shows mean and 95% CI. Statistics performed with one-way ANOVA. (***) P < 0.001; (****) P < 0.0001.

We observed that these protein fragments were sufficient to target proteins to tau aggregates proportionate to their serine content. Specifically, Frag_2, with two polyserine domains, accumulated robustly in cytoplasmic tau aggregates; Frag_1, with one polyserine domain, accumulated to a lesser extent; and neither Frag_0 nor Halo alone, which lack polyserine domains, were enriched in tau aggregates (Figure 2B-C). Similarly, the serine-rich region of PNN was sufficient to target Halo to tau aggregates (Figure 2E-F).

To determine whether polyserine itself is sufficient for localization to tau aggregates, we expressed and verified Halo-tagged polyserine repeats of varying lengths (42, 20, 10, 5) (Figure S2B). We then monitored their recruitment to tau aggregates in HEK293 biosensor cells.

We observed that 42 consecutive serine residues are sufficient to robustly target to both cytoplasmic and nuclear tau aggregates (Figure 3A). We also observed significant enrichment of Halo in tau aggregates with 20-serine residues, and little to no enrichment with the 10 or 5-serine residues (Figure 3B). Halo-tagged 42-serine was also robustly recruited to tau aggregates in H4 neuroglioma cells expressing full length 0N4R P301S tau demonstrating this is not unique to HEK293 cells expressing the tau K18 fragment (Figure 3D). Thus, polyserine alone is sufficient to mediate associations with tau aggregate in a length dependent manner. This provides a molecular explanation for the recruitment of the polyserine containing SRRM2 protein to tau aggregates in cell line and mouse models, as well as in patient samples (Lester et al., 2021; McMillan et al., 2021).

**Figure 3:**
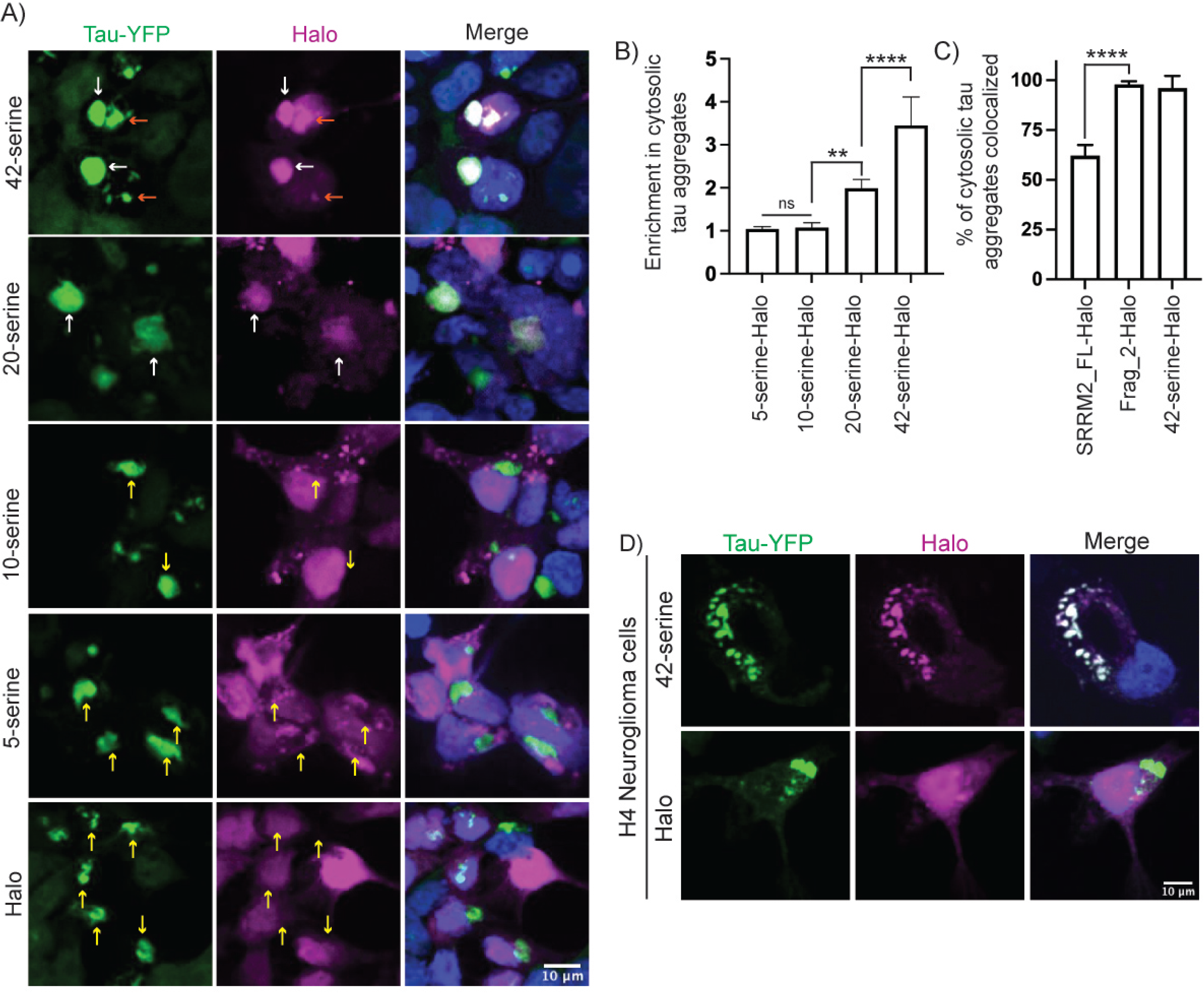
Serine repeats are sufficient for enrichment in tau aggregates. **(A)** Immunofluorescence of tau-YFP (green), Halo (magenta), and DAPI (blue) in HEK293 biosensor cell lines transfected with constructs expressing 42, 20, 10, 5 Serine-Halo, or Halo alone. Enrichment of Halo signal in cytoplasmic tau aggregates is denoted with white arrows, enrichment in nuclear tau aggregates is denoted with orange arrows, and yellow arrows show lack of enrichment. **(B)** Quantification of the ratio of mean intensity of Halo within cytoplasmic tau aggregates relative to Halo signal in the surrounding cytoplasm for serine constructs in (A). n = 40 tau aggregates. Data represent mean and 95% CI. Statistics performed with one-way ANOVA. (**) P = 0.001; (****) P < 0.0001. **(C)** Comparison of the percentage of tau aggregates with colocalization between SRRM2_FL-Halo, Frag_2-Halo, and 42-polyserine-Halo. Data represent mean and 95% CI. n=10. Statistics performed with one-way ANOVA. (****) P < 0.0001. **(D)** Immunofluorescence of tau-YFP (green), Halo (magenta), and DAPI (blue) in H4 neuroglioma cells expressing 0N4R P301S tau transfected with Halo and 42-serine-Halo. Cells were fixed and images 48 hours post seeding.

Interestingly we also observed that 42, 20, 10, and 5-serine-Halo produced cytoplasmic foci that were not present in Halo alone (Figure 3A), suggesting polyserine has self-assembly properties (see below).

### Cytoplasmic SRRM2 and PNN containing assemblies are preferential sites for tau seed propagation into aggregates

To examine the temporal mechanisms through which SRRM2 co-localizes with cytoplasmic tau aggregates, we performed live imaging of HEK293 tau biosensor cells expressing endogenous SRRM2-Halo or PNN-Halo fusion proteins following seeding with tau transgenic mouse brain extracts. These experiments revealed the following key observations.

First, we observed that both SRRM2 and PNN can form two related transient cytoplasmic condensates. The first are MIGs, which contain nuclear speckle proteins, and form during mitosis following nuclear envelope breakdown (Figure 4A) (Prasanth et al., 2003; Rai et al., 2018). MIGs behave like typical condensates exhibiting a round shape and undergoing rapid fusion (Movie 1). We also observed SRRM2 and PNN stochastically form cytoplasmic assemblies independent of mitosis (Figure 4A, Movie 2, 3, 6). Similar cytoplasmic assemblies of SRRM2 have been previously observed in tissue culture cells and human neurons (Berchtold et al., 2018; Tanaka et al., 2018). Since these assemblies contain SRRM2 and PNN (Figure 4A), which are typically found in nuclear speckles, we refer to these assemblies as cytoplasmic speckles (CS).

**Figure 4:**
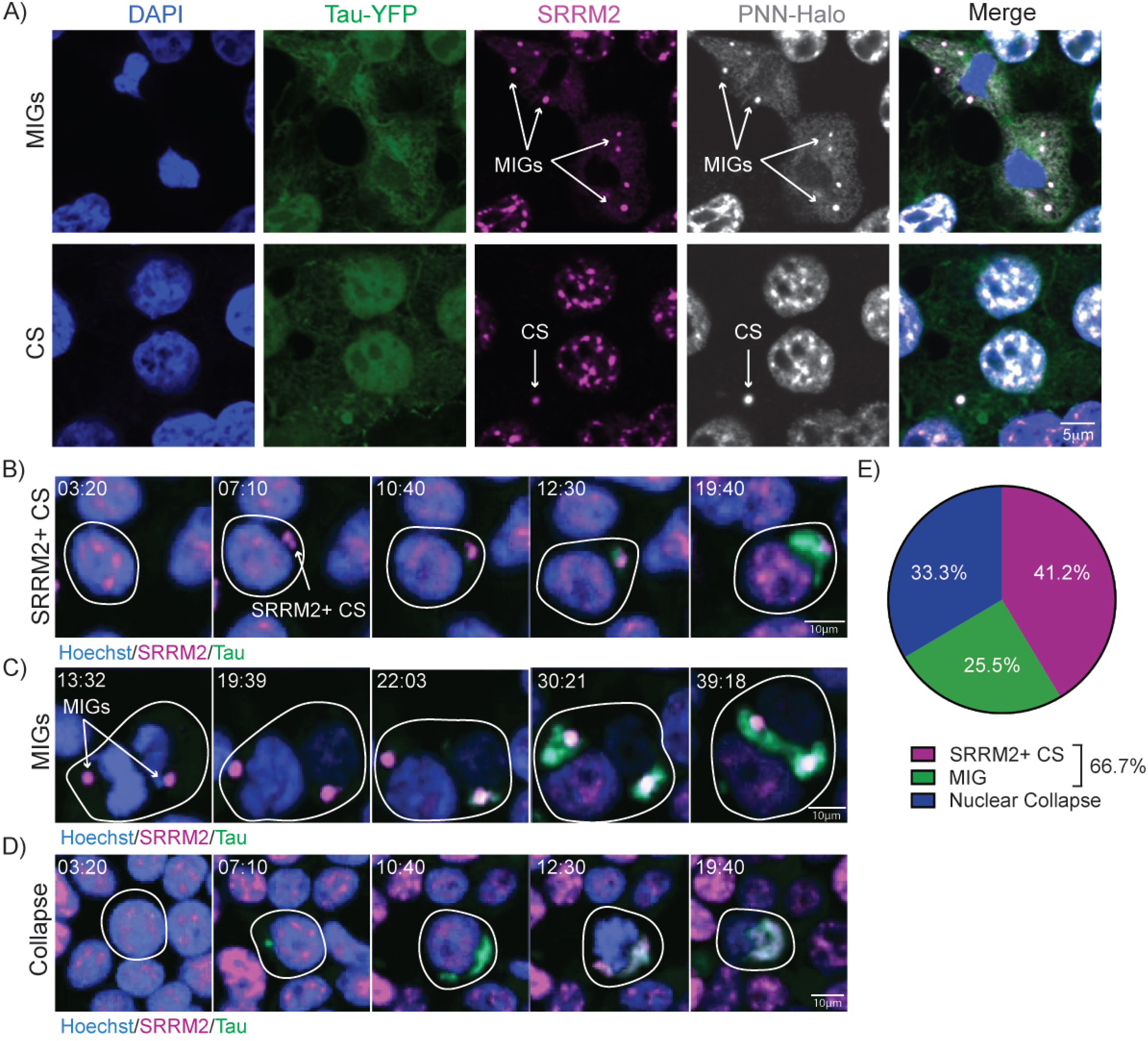
Endogenous SRRM2 assemblies associate with cytoplasmic tau aggregates. **(A)** Immunofluorescence of DAPI (blue), Tau-YFP (green), SRRM2 (magenta) and PNN-Halo (gray) showing the two types of SRRM2+ and PNN+ cytoplasmic assemblies: mitotic interchromatin granules (MIGs) and cytoplasmic speckles (CSs). MIGs are associated with cell division (defined by breakdown of nuclear membrane and chromatin condensation) and CSs are not associated with cell division (no evidence of nuclear membrane breakdown or chromatin condensation). **(B-D)** Live imaging of Hoechst (blue), Tau-YFP (green) and SRRM2_FL-Halo (red) in HEK293 tau biosensor cells seeded with tau aggregates and monitored for 48 hours in 10-minute increments. Stills from live imaging display tau aggregate formation at SRRM2 CSs (A) (Movie 3), MIGs (B) (Movie 4), and aggregate formation followed by nuclear collapse (C) (Movie 5). Time since the onset of imaging is displayed. **(E)** Quantification of the incidence of each mechanism from (A-C). 51 tau aggregates containing SRRM2 at the end of the movie were identified and then aggregates were scored by which mechanism led to the incorporation of SRRM2.

Second, we observed two methods through which SRRM2 colocalized with cytosolic tau aggregates, which also illustrated how tau seeds propagate into larger aggregates in cells. In some cells, we observed growth of tau aggregates that initiated at SRRM2+ CSs (Figure 4B, Movie 3) or SRRM2+ MIGs (Figure 4C, Movie 4). We quantified the percentage of tau aggregates that contained SRRM2 at the end of the video and found that 41.2% of aggregates began in close proximity (<2uM) to CSs and 25.5% began in close proximity to MIGs. Collectively, these two mechanisms represent 66% of the SRRM2+ tau aggregates (Figure 4E).

In the other cells, tau aggregates formed in the cytoplasm independently of any visible SRRM2+ condensate, followed by a rapid (<10 minute) bulk movement of SRRM2 from the nucleus to the cytoplasmic tau aggregate, which we refer to as nuclear collapse (Figure 4D, Movie 5, 33.3%). Whether these tau aggregates initiate on smaller SRRM2+ assemblies below the detection of light microscopy or in association with other subcellular structures remains to be assessed.

We also observed similar results from live imaging of HEK293 tau biosensor cells with endogenously labeled PNN where either tau aggregation initiates at pre-existing PNN+ CSs or MIGs and remains associated while the tau aggregates are growing (Figure S3A, B; Movie 6,7). PNN is also recruited to existing aggregates following nuclear collapse (Figure S4C; Movie 8).

These results identify MIGs and CSs as subcellular assemblies that are preferred sites for the propagation of tau aggregates, with this mechanism occurring in over half of all cases where tau aggregates contain SRRM2 (Figure 4E). Strikingly, in cells that contain observable MIGs or CSs, we typically observe the initial growth of the tau aggregate occurring in conjunction with the MIG or CS. MIGs contain snRNAs (Ferreira et al., 1994), which is notable since tau aggregates in patients can contain U1 snRNAs (Hales et al., 2014a, 2014b), and tau aggregates in cell lines and mouse models are enriched in snRNAs (Lester et al., 2021). This provides a possible explanation for why snRNAs and nuclear RNA binding proteins accumulate in tau aggregates in disease.

### Stress granules are not preferential sites for tau seed propagation into aggregates

Tau aggregates could preferentially form in association with MIGs and CSs due to their inclusion of RNA, or due to other protein domains within these assemblies – such as polyserine domains – that may affect tau propagation. To test if another cytoplasmic RNP assembly can preferentially propagate tau seeds, we examined if tau aggregates similarly formed in association with stress granules. Stress granules are assemblies of untranslating mRNPs that form when translation initiation is inhibited (Protter and Parker, 2016), and have been proposed to associate with pathological inclusions in neurodegenerative disease (Cruz et al., 2019). We modified HEK293 biosensor cells to stably express mRuby or mRuby-tagged G3BP1 – a canonical stress granule marker. To induce stress granules that would persist during the course of tau aggregation, we treated cells with Pateamine A (PatA), which inhibits translation initiation by disrupting eIF4A function (Dang et al., 2006).

Following the addition of tau seeds and 50nM PatA, we observed stress granules and tau aggregates were mostly independent, with limited overlap or docking of mRuby-G3BP1 with tau aggregates at a late timepoint (Figure S4A, B). Live imaging of tau aggregate formation under conditions of PatA-mediated stress granule induction demonstrated tau aggregates formed independently of stress granules but could subsequently exhibit transient surface docking with stress granules (Figure S4C, Movie 9,10). Prior studies have shown tau associates in model systems and patient samples with TIA1, a nuclear protein that relocalizes to cytoplasmic stress granules under stress but – consistent with our findings – does not colocalize with other stress granule markers, most notably G3BP1 (Apicco et al., 2018; Vanderweyde et al., 2012). Collectively, these results highlight that the engagement of tau – and in the case of MIGs and CSs the preferential propagation of tau fibers – with RNP granules is not a general property, but rather is specified by unique features of these assemblies.

### Cytoplasmic assemblies formed by exogenous polyserine-containing proteins are sufficient to create conducive sites for tau aggregation

Since polyserine domains are sufficient to interact with tau aggregates, and are enriched in proteins found in nuclear speckles, MIGs, and CSs, we hypothesized that polyserine containing protein domains, or polyserine itself, might be the driving principle for the formation of assemblies that serve as a preferred sites for tau propagation. Consistent with this hypothesis, we observed that overexpression of 42-serine and SRRM2-Frag_2, but not Halo alone, led to the formation of cytoplasmic assemblies (Figure 5A-C).

**Figure 5:**
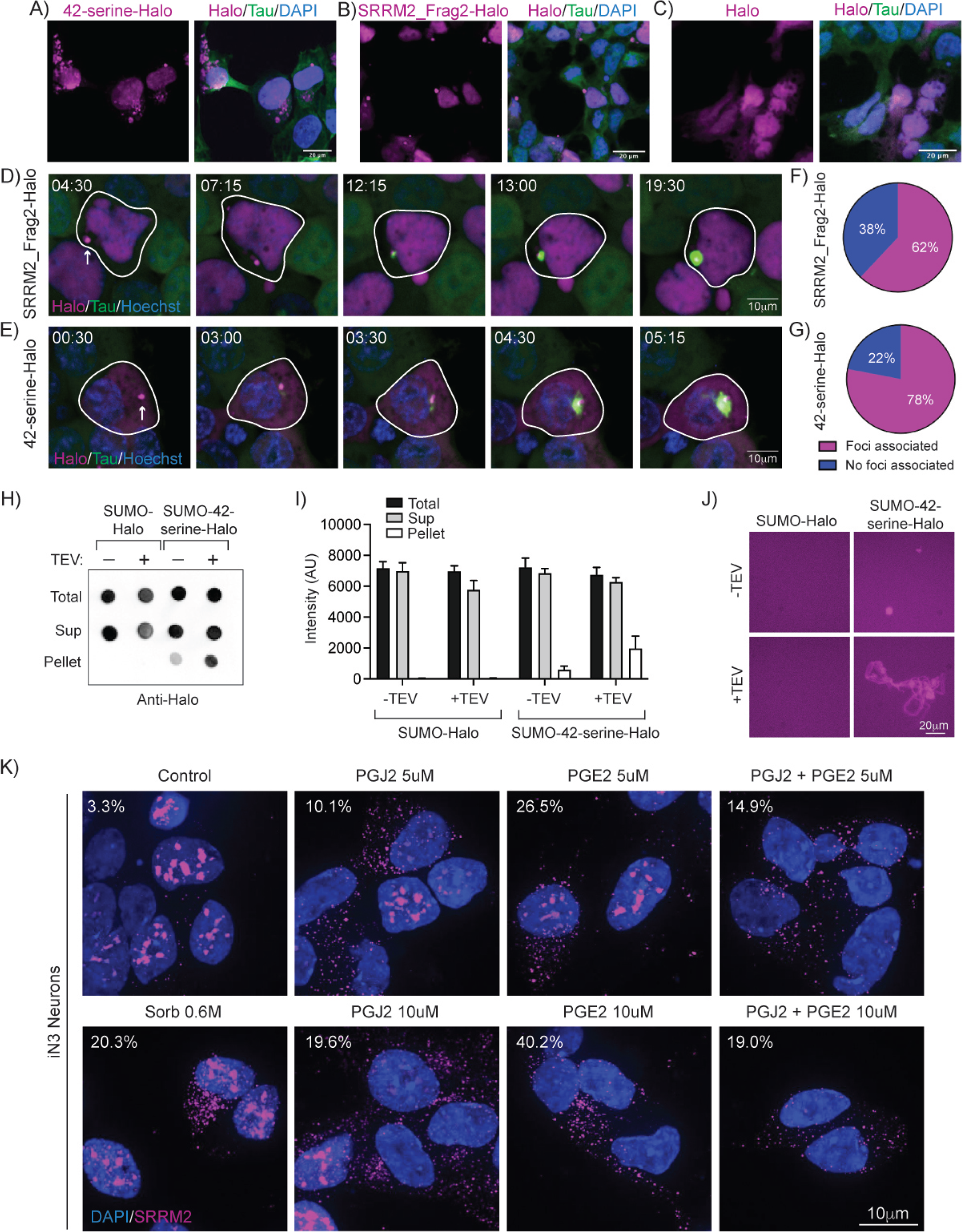
Polyserine-based assemblies are sites of tau aggregate formation. **(A, B, C)** Immunofluorescence of Halo (magenta), Tau-YFP (green) and DAPI (blue) in HEK293 biosensor cells transfected with 42-serine-Halo, SRRM2_Frag2-Halo, or Halo. **(D)** Stills from live-imaging of HEK293 tau biosensor cells expressing Tau-YFP (green), transfected with SRRM2_Frag2-Halo (magenta) and labeled with Hoechst (blue). Cells were lipofected with clarified tau brain homogenate and imaged for 24 hours at a time interval of 15 minutes. Time since the onset of imaging is displayed. (Movie 11). **(E)** Stills from live-imaging of HEK293 tau biosensor cells expressing Tau-YFP (green), transfected with 42-serine-Halo (magenta) and labeled with Hoechst (blue). Cells were lipofected with clarified tau brain homogenate and imaged for 24 hours at a time interval of 15 minutes. Time since the onset of imaging is displayed. (Movie 12). **(F-G)** Quantification of 50 tau aggregates that contained Halo signal at the termination of live imaging scored by whether tau aggregation initiated within 2uM of cytoplasmic Halo+ foci in cells transfected with SRRM2_Frag2-Halo (F) or 42-serine-Halo (G). **(H)** Dot blot probed with anti-Halo antibody of total, supernatant (Sup) and pellet from recombinant SUMO-Halo and SUMO-42-serine-Halo preparations with or without TEV protease cleavage. **(I)** Quantification of dot blot from groups as in (H). n=5. Data represent mean and SEM. **(J)** Imaging of samples as in (H) labeled with Halo ligand (magenta). **(K)** Immunofluorescence of SRRM2 (magenta) with DAPI (blue) stain in human iPSC derived cortical neurons (iN3 neurons) at 12 days post-differentiation treated with either vehicle control, 0.6M sorbitol for 1 hour, PGJ2 for 15 hours (5uM or 10uM), and/or PGE2 for 15 hours (5uM or 10uM). Percentages in upper left-hand corner show the percentage of neurons that have MIGs after being treated with the corresponding compound.

Through live cell imaging, we observed tau aggregates preferentially formed in association with both SRRM2-Frag_2-Halo and 42-serine-Halo assemblies, similar to endogenously labeled assemblies of SRRM2 and PNN (Figure 5D,E, Movies 11,12). In transfected cells expressing SRRM2-Frag_2-Halo or 42-serine-Halo that had Halo+ assemblies, 62% and 78% of tau aggregates initiated near those Halo+ assemblies (<2uM), respectively (Figure 5F,G). Thus, polyserine is sufficient to create assemblies that establish a local environment conducive for tau aggregation.

Because 42-serine-Halo forms assemblies in cells, we next evaluated whether 42-serine-Halo forms homotypic assemblies in vitro. We observed that purified recombinant SUMO-42-serine-Halo formed visible assemblies that pelleted at 100,000 xg, while SUMO-Halo alone did not (Figure 5H, I). Moreover, treatment of SUMO-42-serine-Halo with TEV to remove the SUMO solubility tag led to an increase in the formation of pelletable assemblies for 42-serine-Halo but not the SUMO-Halo (Figure 5H, I). Microscopic examination of the assemblies showed that SUMO-42-serine-Halo formed visible assemblies that ranged in size from 2.74 µM to 27 µm with an average size of 10.3 µM. Removal of the SUMO tag by TEV cleavage led to the formation of larger 42-serine-Halo assemblies that ranged in size from 2.79 µM to 199 µM with an average size of 45.8 µM. We did not observe any visible assemblies with SUMO-Halo even after removal of the SUMO tag (Figure 5J). Taken together, our *in vitro* analysis of 42-serine supports our *in vivo* observations that polyserine can self-assemble.

### Stress induces cytoplasmic speckles in iPSC neurons

The results above suggest that the formation of CSs in post-mitotic neurons might create condensates that would enhance the propagation of tau aggregates. Previous results have shown that amyloid-β toxicity can induce cytoplasmic assemblies of SRRM2, which we infer to be CSs, in mouse and human neurons (Tanaka et al., 2018). To determine if other stresses can induce CSs in post-mitotic neurons, we treated induced pluripotent stem cell (iPSC) derived cortical neurons with various stressors (Fernandopulle et al., 2018; Wang et al., 2017). We validated the neuronal identity of the iPSC derived cortical neurons using two neuron specific markers, β-III tubulin and MAPT (Figure S5A-C) (Fernandopulle et al., 2018). Strikingly, we observed SRRM2+ CSs in neurons following hyperosmotic sorbitol stress, as well as inflammatory stress induced through treatment with prostaglandin J2 (PGJ2) and prostaglandin E2 (PGE2) (Figure 4K, S5D), which are elevated in chronic inflammatory states and neurodegeneration (Figueiredo-Pereira et al., 2016; Tauber and Parker, 2019; Xia et al., 2015). These results show external triggers, including aspects of neuroinflammation, can promote the formation of SRRM2+ CSs in neurons.

### Tau aggregate formation is modulated by levels of polyserine containing proteins

The results above suggest cytoplasmic assemblies enriched in polyserine domains form biochemical environments conducive to tau aggregation, which predicts that the level of polyserine repeat domains would correlate with tau aggregate formation. To examine this possibility, we increased or decreased the amount of polyserine regions in cells and examined tau aggregate formation in response to seeding. To quantify tau aggregation, we utilized FRET based flow cytometry of the HEK293T tau biosensor cells as a measure of tau aggregation (Furman et al., 2015; Holmes et al., 2014). Gating parameters were performed as previously reported (Figure S6A-C) (Holmes et al., 2014), and tau seeding leads to a shift of cells into the FRET+ population (Figure S6C, D). We validated this assay by showing knockdown of MSUT2 (mammalian suppressor of tauopathy 2) – a nuclear speckle protein shown to alter tau aggregation in mouse and invertebrate model organisms (Wheeler et al., 2019) – led to a reduction in both the percentage of tau aggregate positive (FRET+) cells and the integrated FRET density (the product of FRET+ percentage and the median fluorescence intensity; FRET density is a combinatorial measure of the aggregation within each cell and a population based analysis of the extent of tau aggregation) (Guthrie et al., 2011; Wheeler et al., 2019) (Figure 6A-C, S6G).

**Figure 6:**
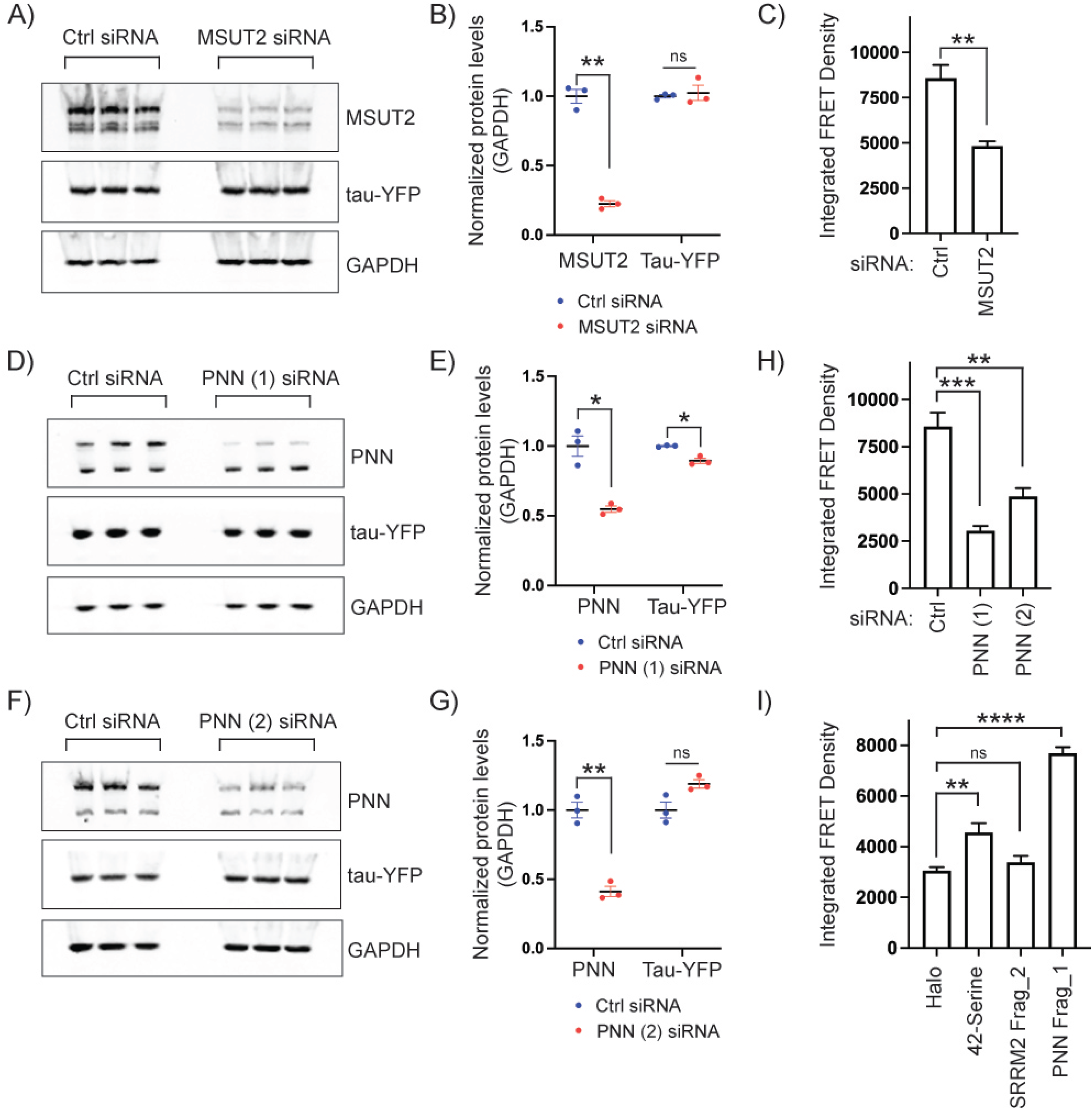
Cellular levels of polyserine containing proteins modulates tau aggregation. **(A)** Western blot of MSUT2, tau-YFP and GAPDH protein levels in HEK293 biosensor cells treated with control or MSUT2 siRNA. **(B)** Quantification of Western blot shown in (A) normalized to GAPDH. Bars represent mean and SEM. Statistics performed with unpaired t-test with Welch’s correction. (ns) > 0.05; (**) P < 0.01. **(C)** Integrated FRET density (product of FRET+ percentage and median fluorescence intensity) of HEK293 biosensor cells treated with control or MSUT2 siRNA measured by flow cytometry and analyzed as in Figure S6A-C. Data represent mean and SEM. Statistics performed with Mann-Whitney test. (**) P < 0.01. **(D)** Western blot of PNN, tau-YFP and GAPDH protein levels in HEK293 biosensor cells treated with control or PNN (1) siRNA. **(E)** Quantification of Western blot shown in (D) normalized to GAPDH. Bars represent mean and SEM. Statistics performed with unpaired t-test with Welch’s correction. (*) P < 0.05. **(F)** Western blot of PNN, tau-YFP and GAPDH protein levels in HEK293 biosensor cells treated with control or PNN (2) siRNA. **(G)** Quantification of Western blot shown in (F) normalized to GAPDH. Bars represent mean and SEM. Statistics performed with unpaired t-test with Welch’s correction. (ns) > 0.05; (**) P < 0.01**. (H)** Integrated FRET density in HEK293 biosensor cells treated with control or each PNN siRNA measured by flow cytometry and analyzed as in (A-F). Data represent mean and SEM. Statistics performed with one-way ANOVA. (**) P < 0.01; (***) P < 0.001. **(I)** Integrated FRET Density in HEK293 biosensor cells transfected with Halo, 42-Serine, SRRM2 Frag_2 and PNN Frag_1 constructs measured by flow cytometry and analyzed by single cell gating in Figure S6A,B, subsequently 561nm Halo+ gating in Figure S7A,B, and lastly for FRET positivity as in Figure S6C. Data represent mean and SEM. Statistics performed with one-way ANOVA. (ns) P > 0.05; (**) P < 0.01; (****) P < 0.0001.

To reduce polyserine containing domains in cells, we used siRNAs to knockdown either SRRM2 or PNN. Successful knockdown of SRRM2 – without alteration in tau protein levels – showed no significant alteration in tau aggregation, as assessed by the percent of FRET positive cells or integrated FRET density (Figure S6I-L). However, since PNN (present at ~1.2 million molecules/cell) is more abundant than SRRM2 (present at ~500,000 molecules/cell) (Itzhak et al., 2016), we hypothesized that SRRM2 knockdown may not sufficiently reduce levels of polyserine. To knockdown PNN, we validated two siRNAs and monitored effects on tau levels (Figure 6D-G). While PNN (1) siRNA led to a modest reduction in tau levels, PNN (2) siRNA resulted in a slight increase (Figure 6D-E,F-G). Regardless of these effects on tau protein levels, knockdown with either siRNA led to a significant reduction in the percentage of FRET+ cells and integrated FRET density (Figure 6H). Thus, a reduction of PNN levels reduces tau aggregation.

To determine whether increasing polyserine concentration affects tau aggregation, we transfected cells with constructs expressing Halo-tagged 42-polyserine, SRRM2-Frag_2, and PNN-Frag_1 and monitored tau aggregation via flow cytometry. An additional gating step to sort for transfected cells based on Halo expression was performed (Figure S7A,B). We found overexpression of all three of these proteins resulted in an upward trend in FRET+ percentage with 42-serine and PNN-Frag_1 reaching statistical significance (Figure S7C-G). Further, 42-serine and PNN-Frag_1 also led to a significant increase in integrated FRET density (Figure 6I). Importantly, expression of each construct did not lead to meaningful increases in tau levels as shown by the distribution of CFP signal in sorted cell populations (Figure S7H-K).

Taken together, these results indicate that decreasing or increasing polyserine domain levels in the cells can correspondingly decrease or increase tau aggregation. This provides evidence that polyserine containing assemblies are not only sites of preferential tau fiber growth, but their level can affect the degree of tau aggregation.

## DISCUSSION

Our observations demonstrate polyserine protein domains mediate interactions with tau aggregates. Polyserine rich domains in the nuclear splicing speckle proteins SRRM2 and PNN are necessary and sufficient for localization to tau aggregates (Figures 1-2). Further, polyserine repeats alone are sufficient for targeting to tau aggregates in a length dependent manner (Figure 3). The association of polyserine with tau aggregates provides a molecular explanation for the mislocalization of SRRM2 to pathogenic tau inclusions in post-mortem samples from AD, CBD, and FTLD patients (Lester et al., 2021; McMillan et al., 2021).

Additional evidence demonstrates that polyserine domains are sufficient to create biological assemblies that are preferential sites of tau fiber propagation. First, we observed that in a seeding model, 66% of tau aggregates that are SRRM2+ initiate in association with either MIGs or CSs (Figure 4). Second, over-expression of either the SRRM2 C-terminal domain containing polyserine runs (SRRM2-Frag_2-Halo), or 5, 10, 20, or 42-polyserine, drives formation of assemblies in cells and similar results can be seen *in vitro* for 42-polyserine, demonstrating polyserine has self-assembly properties (Figure 5). Third, the condensates formed by over-expression of 42-polyserine can also serve as preferred sites of tau aggregation (Figure 5). Fourth, knockdown of PNN leads to a reduction in the formation of tau aggregates (Figure 6). Finally, we observed overexpression of 42-polyserine or the PNN C-terminus (PNN-Frag_1-Halo) increased tau aggregation (Figure 6). A unifying model is that polyserine domains can self-assemble and define a biochemical environment that can promote tau fiber growth.

The observation that MIGs, CSs, or artificial polyserine-based cytoplasmic assemblies serve as preferred sites of tau fiber propagation provides an explanation for nuclear speckles – which are rich in polyserine containing proteins (Supplemental Table 1) – serving as a second preferred site of tau fiber propagation (Lester et al., 2021). Moreover, a 42-serine repeat is sufficient to target exogenous proteins to nuclear tau aggregates (Figure 3A, orange arrows), demonstrating the molecular interactions between polyserine and tau are similar in both cytoplasmic and nuclear tau aggregates.

This work illustrates the fundamental principle that growth of aberrant protein fibers can be enhanced by specific subcellular locations with defined biochemical composition. Our results indicate the effects of polyserine containing assemblies are not a general principle of RNP assemblies as G3BP1+ stress granules do not associate with tau aggregates during formation (Figure S4). In principle, a polyserine defined biochemical environment could enhance the rate of tau fiber growth in several, possibly overlapping, manners. Through interaction with tau monomers, it could increase the local concentration of tau and the probability of tau-tau interactions. One attractive possibility is that polyserine is sufficient to nucleate a condensate that recruits additional factors and tau to promote fibrillization as has been seen with other fibrilization prone proteins (Lin et al., 2015; Molliex et al., 2015; Patel et al., 2015). Alternatively, polyserine interaction with tau could stabilize a particular fold of either the tau monomer or seed that enhances fiber growth. For example, since polyserine can form a type of fiber related to a coiled-coil (Lilliu et al., 2018), it could provide a template for the increased rate of tau fiber growth. Another possibility is that polyserine, or a polyserine-interacting molecule, could serve as a co-factor for tau propagation. Interestingly, since polyserine domains can be heavily phosphorylated (Cesaro and Pinna, 2015), a phosphorylated polyserine region could function similarly to other polyanions such as RNA or heparin that promote tau fibrillization in vitro (Dinkel et al., 2015; Friedhoff et al., 1998; Kampers et al., 1996). Further work will be necessary to elucidate the mechanisms through which polyserine regulates tau aggregation.

Multiple observations suggest that the interaction between polyserine domains and tau will be pertinent to human disease. First, SRRM2 is known to mislocalize to tau aggregates in post-mortem samples from AD, CBD, and FTLD patients (Lester et al., 2021), and the degree of mislocalization corresponds with increased severity of pathological tau deposition in humans and mouse models (McMillan et al., 2021). Second, previous work has shown that β-amyloid deposition can promote SRRM2 phosphorylation and export to the cytoplasm leading to disruptions in neuronal RNA splicing (Tanaka et al., 2018). Third, we observed SRRM2+ CSs form in non-dividing human iPSC derived cortical neurons when stressed with the inflammatory compounds PGJ2 and PGE2 as well as sorbitol induced hyperosmolar stress (Figure 5). This suggests that β-amyloid abnormalities, or other inflammatory triggers, could promote the relocalization of SRRM2 and PNN to the cytoplasm, promote CS formation, and lead to increased tau fibril growth. This could provide a molecular link explaining how β-amyloid plaques and other inflammatory compounds can increase the probability of tau tangle formation in human AD.

The polyserine-mediated interactions of tau with nuclear speckle components, either in the nucleus or the cytosol may contribute to neurotoxicity through disrupting RNA homeostasis. One possible mechanism of neurotoxicity is sequestration of SRRM2 or PNN in cytoplasmic tau aggregates leading to loss of their physiologic function. SRRM2 is highly conserved throughout evolution and loss of function mutations in SRRM2 have been shown to cause neurodevelopmental disorders suggesting that perturbations leading to SRRM2 loss of function are deleterious to neurons (Cuinat et al., 2022). Additional evidence suggests that loss of SRRM2’s nuclear function leads to splicing abnormalities through proteins such as PQBP1 (Tanaka et al., 2018). Furthermore, deletion of the nuclear speckle protein MSUT2 (Guthrie et al., 2011; Lester et al., 2021) has been shown to suppress tau toxicity in mouse models and decreases tau aggregates in cells (Wheeler et al., 2019) (Figure 6) further connecting tau toxicity to nuclear speckle components.

Taken together, we suggest a working model for the relationship between tau aggregation and RNP granules with the following key principles (Figure 7). First, when tau seeds form or enter a cell, they can be cleared, or propagate new fibers with or without an association with MIGs and CSs. We hypothesize that the rate of tau fibrillization and/or propagation will be increased in neurons with SRRM2+ and/or PNN+ CSs. Conditions that increase CSs, such as the presence of high concentrations of neuroinflammatory compounds like PGE2 and PGJ2, or proximity to amyloid-β plaques (Tanaka et al., 2018) could increase tau propagation in neurons. An important continuation of this study will be to determine what additional extracellular or intracellular events increase CS formation in post-mitotic neurons and assess how those events influence tau propagation and downstream mechanisms of neurodegeneration.

**Figure 7:**
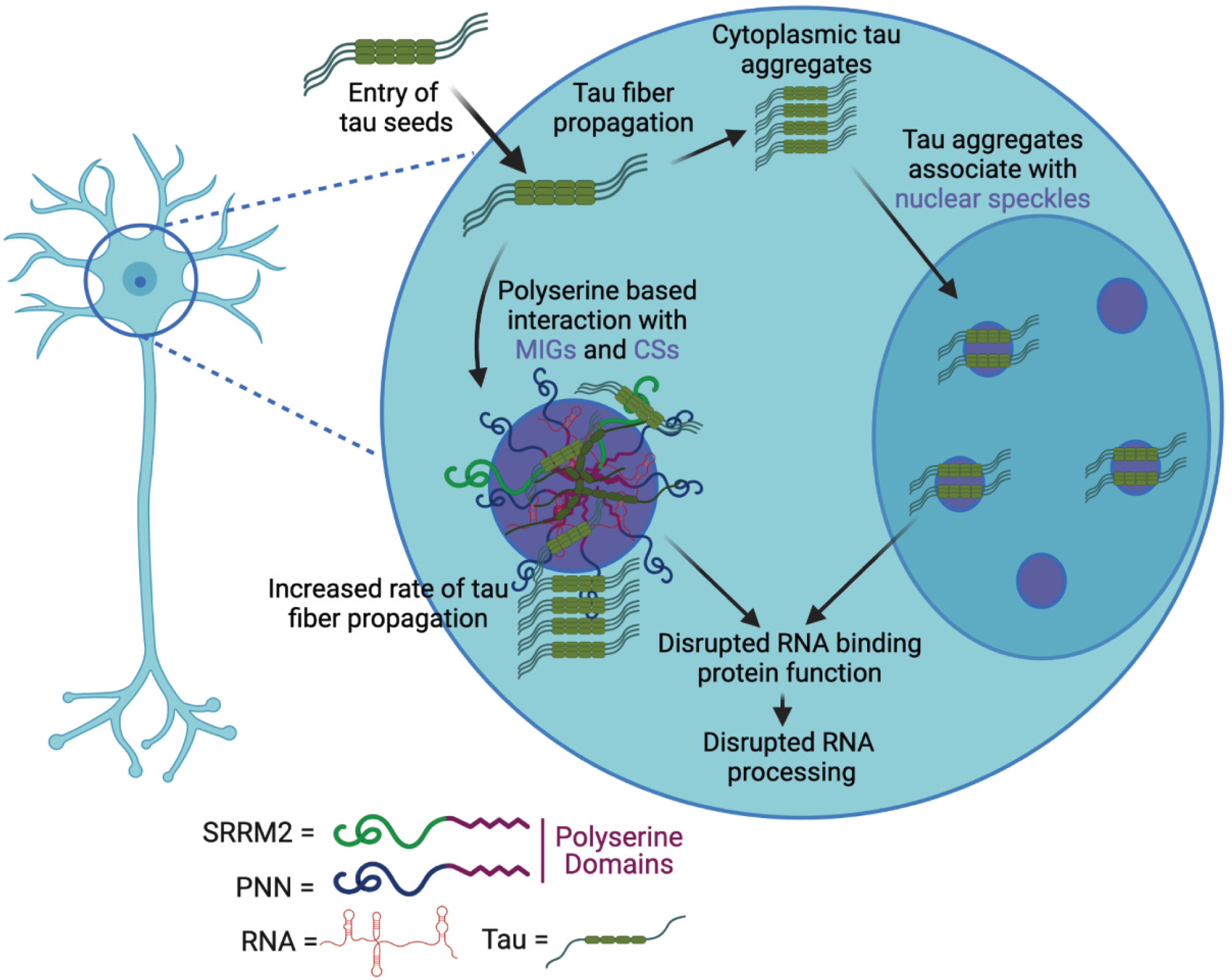
Model. **(A)** Model showing a possible mechanism by which tau seeds introduced into HEK293 tau biosensor cells can form cytoplasmic tau aggregates or be recruited to SRRM2+ and PNN+ cytoplasmic speckles (CSs) or mitotic interchromatin granules (MIGs) that contain polyserine domains and can be derived from nuclear speckles. Tau aggregates can grow and propagate within MIGs and CS. Tau’s interaction with RNA binding proteins such as SRRM2 and PNN in MIGs, CSs, and nuclear speckles could induce a loss of function in RNA binding protein function and lead to disruptions in RNA processing.

## MATERIALS AND METHODS

### Cell culture and tau aggregate seeding of HEK293 cells

As previously described (Lester et al., 2021; Sanders et al., 2014), HEK293 biosensor cells stably expressing the 4R repeat domain of tau (K18) with the P301S mutation were purchased from ATCC (CRL-3275) (previously described in (Holmes et al., 2014)). Cells were seeded at 1.25 x 10^5^ cells/mL in 500uL of DMEM with 10% FBS and 0.2% penicillin-streptomycin antibiotics on PDL coated glass coverslips in a 24-well tissue culture treated plate (Corning 3526) and allowed to grow overnight in incubators set to 37°C with 5% carbon dioxide. The next day, 7ug of 1 mg/mL clarified P301S tau mouse brain homogenate or 7uL PBS was mixed with 6uL of Lipofectamine 2000 and brought up to 100uL in PBS and allowed to sit at room temperature for 1.5 hours. The mixture was then added to 300uL of DMEM without FBS or antibiotics and mixed by pipetting. 50uL of this mixture was added to each well of a 24 well plate and allowed to incubate at 37 °C for 24 hours. Tau aggregate formation was monitored using a fluorescence microscope with a 488nm filter.

### Cell culture and tau seeding of H4 biosensor cells

As previously described (Lester et al., 2021), H4 cells (ATCC Cat# HTB-148, RRID:CVCL_1239) stably expressing the pIRESpuro3 vector (Clontech) containing a codon-optimized 0N4R MAPT gene with the P301S point mutation and tagged with YFP were cultured in Dulbecco’s Modified Eagle’s Medium (DMEM) supplemented with 10% fetal bovine serum (FBS) and 0.2% penicillin-streptomycin, and maintained in incubators set to 37°C with 5% carbon dioxide. Cells were plated in a 12-well glass-bottomed dish at 1.25×10^5^ cells/well and allowed to settle for a minimum of 2 hours prior to infection with tau seeds (described above). Cells were grown for 48 hours in the presence of tau seeds prior to fixation.

### Clarification of brain homogenate for tau aggregate seeding in HEK293 cells

As previously described (Lester et al., 2021), 10% brain homogenate from Tg2541 or WT mice was centrifuged at 500 x g for 5 minutes, the supernatant was transferred to a new tube and centrifuged again at 1,000 x g for 5 minutes. The supernatant was again transferred to a new tube and the protein concentration was measured using bicinchoninic acid assay (BCA), and diluted in DPBS to 1 mg/mL for transfection into HEK293 tau biosensor cells.

### Immunofluorescence

As described above, cells were grown in 24 well plates on PDL coated coverslips, fixed for 10 minutes in 4% paraformaldehyde, washed 3x with PBS, and permeabilized with 0.1% Triton-X100 for 10 minutes. Cells were then washed with PBS 3X, blocked in 5% BSA for 30 minutes, followed by addition of primary antibodies (Supplemental Table 2) at indicated concentration in 5% BSA and incubated overnight at 4 deg on a rotator. Cells were washed 3X with PBS and incubated with secondary antibody in 5% BSA for 30 minutes at room temperature on a rotator. Cells were then washed 3X with PBS and incubated in DAPI for 5 minutes at a final concentration of 1μg/mL in PBS. Cells were washed one more time with PBS and mounted using ProLong Glass Antifade Mountant.

### Generation of cell lines

For generation of CRISPaint edited cell lines, HEK293 tau biosensor cells were seeded at 5*10^5^ cells/ml in 6 well plates and allowed to grow overnight in a 37°C incubator. The next day, Lipofectamine3000 was used to transfect cells with 0.5ug of pSpCas9(BB)-2A-GFP(PX458) targeting plasmid (Addgene # 48138) containing sgRNA sequences targeting full-length or truncations in SRRM2 and PNN (Supplemental Table 3), 0.5ug of the respective pCAS9-mCherry-Frame selector plasmid (Addgene #66939, 66940, 66941), and 1μg of pCRISPaint-HaloTag-PuroR plasmid (Addgene #80960). After 24 hours, cells were selected with 2ug/mL puromycin. JF646 was added to the media at a final concentration of 200nM for 24 hours to covalently label the Halo fusion proteins for visualization by fluorescence imaging and analysis by gel electrophoresis.

For generation of mRuby-tagged cell lines, constructs encoding mRuby2 or mRuby2-G3BP1 in a pLenti EF1 vector backbone were cloned. HEK293T WT cells were transfected using Lipofectamine3000 with pLenti plasmids and lentiviral packaging plasmids (Gag-pol (Addgene #14887), VSV-G (Addgene #8454), rSV-Rev (Addgene #12253)). Lentivirus was used to transduce HEK293 tau biosensor cells followed by selection with blasticidin (2μg/mL).

### Cloning and expression of SRRM2 and PNN C-terminal fragments and polyserine repeats

To express SRRM2 C-terminal fragments in a pcDNA-PuroR expression plasmid, RNA was extracted from SRRM2_FL-Halo cells using Trizol, and reverse transcribed to cDNA using oligodT primers and Superscript III. Regions in the C-terminus of SRRM2_FL-Halo were amplified by PCR and cloned into EcoRV/XbaI digested pcDNA plasmid using In-Fusion cloning. For polyserine-Halo and PNN_Frag1 constructs, gene blocks were ordered from IDT with codons optimized for synthesis then cloned via In-Fusion into the pcDNA plasmid backbone.

### Gel electrophoresis and Western blotting

For analysis of SRRM2 truncation cell lines, cells were grown in a 6 well plate to 50-80% confluence, washed 1x with PBS, and trypsinized in 0.5mL of trypsin. Cells were collected in a 1.5mL microcentrifuge tube and centrifuged at 500g for 5 minutes, washed 1x with PBS, and brought up in 100uL of lysis buffer (25mM Tris pH 7.5, 5% glycerol, 150mM NaCl, 2.5mM MgCl_2_, 1% NP-40, 1:20 BME, 1X phosphatase/protease inhibitor). Lysate was pipetted up and down to mix and incubated on ice for 5 minutes. Lysate was then centrifuged at 16,000g for 5 minutes and supernatant was transferred to a new tube and protein concentration was measured via Bradford. 10-15 ug of protein was combined with 4X LDS loading dye and boiled for 7 minutes prior to loading on a NuPAGE 4 to 12% Bis-Tris mini protein gel. Gels were directly imaged to examine covalently linked JF646.

For validation of PNN CRISPaint edited cell lines, 6×10^5^ cells were plated in a 6-well format 24 hours before addition of JF646 was added at a final concentration of 200uM and collected 24 hours after labeling. Cell pellets were lysed in 2X SDS loading buffer, passed through a 25G syringe, and boiled. Extracts were run on 4-20% Tris-Glycine protein gel and imaged directly for JF646 fluorescence. The gel was then transferred using iBlot 2 Transfer Device (Thermo Fisher) to nitrocellulose membranes for Western blotting.

For Western blotting, membranes were blocked in 5% milk in Tris-buffered Saline with 0.1% Tween (TBS-T) for 1 hour, incubated with primary antibodies (Supplemental Table 2) in TBS-T for 2 hours at room temperature, washed 3 x 10 minutes with TBS-T, incubated with secondary antibodies in TBS-T for 1 hour at room temperature, then washed 6 x 5 minutes with TBS-T before developing with Clarity Western ECL Substrate (Bio-rad).

### Live cell imaging

For analysis of tau aggregate formation and SRRM2 relocalization, HEK293 tau biosensor cells with SRRM2_FL-Halo were seeded in 24 well glass bottom plates with #1.5 cover glass at 2.5*10^5^ cells/ml with or without 200nM JF646-Halo ligand and allowed to grow overnight at 37°C. To counter stain nuclei, Hoechst 33342 was added to the cell culture media and allowed to incubate for 15 minutes prior to imaging. For analysis of SRRM2 relocalization during tau aggregation, cells were imaged on an Opera Phenix High Content imaging system where images in the Cy5 (Halo-JF646), GFP (Tau-YFP), and DAPI (Hoechst) channels were acquired every 10 minutes for 48 hours at 37°C and 5% CO_2_.

For live imaging of tau aggregate formation and PNN relocalization, HEK293 tau biosensor cells with PNN_FL-Halo labelling were seeded at a density 1.25 x 10^5^ cells per 24-well in poly-L-lysine coated glass bottom plates. The following day, media was changed to Fluorobrite DMEM supplemented with 10% FBS and seeded using Lipofectamine3000 with tau brain homogenate. 5 hours post-seeding JF646 ligand (200nM) and Hoechst 33342 were added. 1 hour post-labeling imaging was started using a Nikon Spinning Disk Confocal acquiring images every 10 minutes for 24 hours.

For live imaging of tau aggregate formation and stress granules, HEK293 tau biosensor cells with mRuby2-G3BP1 were plated and seeded as described for PNN-Halo live imaging. 5 hours post-seeding Hoechst 33342 (1μg/mL) and Pateamine A (50nM) were added and after 30 minutes imaging was initiated with acquisition every 10 minutes for the following 24 hours.

For live imaging of transiently transfected 42-serine and SRRM2_Frag2-Halo constructs and tau aggregate formation, HEK293 tau biosensor cells were seeded at a density of 1.00 x 10^5^ cells per 24-well in poly-L-lysine coated glass bottom plates. The following day, cells were transfected using Lipofectamine3000 with 500ng plasmid per well. 24 hours post-transfection cells were seeded with tau brain homogenate. 5 hours post-seeding JF646 ligand (200nM) and Hoechst 33342 were added. 1-hour post-labeling imaging was started using a Nikon Spinning Disk Confocal acquiring images every 15 minutes for 24 hours.

### Protein expression and purification

A gene block for codon optimized 42-serine-Halo and Halo were purchased from IDT and used to subclone 42-serine-Halo and Halo into pET28a plasmid modified to have an N-terminal 6xHis-SUMO tag. SUMO-Halo and SUMO-42-serine-Halo were expressed in Rosetta2(DE3)pLysS e. coli in LB for 4 hours at 37°C after induction with 200 uM IPTG. The cell pellets were resuspended in lysis buffer (50 mM MOPS pH 7.0, 300 mM NaCl, 0.1% NP-40, 30 mM Imidazole, 1 mM DTT) supplemented with complete ULTRA EDTA-free protease inhibitors and 0.1 mM AEBSF and lysed by sonication. Lysate was labeled with JF549 Halo ligand for 30 min at room temperature. Proteins were purified using Ni2+-NTA resin and eluted with 50 mM MOPS pH 7.0, 300 mM NaCl, 0.1% NP-40, 300 mM Imidazole, and 1 mM DTT supplemented with complete ULTRA EDTA-free protease inhibitors and 0.1 mM AEBSF. Eluted proteins were dialyzed overnight at room temperature into 50 mM MOPS pH 7.0, 100 mM NaCl, 0.1% NP-40, and 1 mM DTT.

### *In vitro* polyserine assembly

To assess assembly formation, 20 µM purified SUMO-Halo and SUMO-42-serine-Halo were incubated with TEV for 22 hours at room temperature in dialysis buffer. Reactions were either subjected to ultracentrifugation, microscopic examination, or boiled in SDS and analyzed by SDS-PAGE. For ultracentrifugation analysis, reactions were spun at 100,000g for 1 hr, washed with dialysis buffer and spun a second time at 100,000g for 30 min. 1 µl samples from the total, supernatant and pellet fractions were spotted onto nitrocellulose and blotted with rabbit anti-HaloTag pAB (Promega). For microscopic examination, reactions were transferred to Bio-One CELLview slides (Greiner) and imaged at 60X magnification on Nikon epifluorescence microscope. For SDS-PAGE analysis, reactions were boiled in 1x SDS-loading dye for 5 min at 95°C, run on a 4-12% Bis-Tris gel, and transferred to nitrocellulose. The membrane was blotted with rabbit anti-HaloTag pAB (Promega).

### Differentiation and Stress of iPSC-derived Cortical Neurons

As previously described (Fernandopulle et al., 2018), WTC-11 cells expressing a doxycycline inducible form of the master neuronal transcriptional regulator neurogenin-2 (NGN2) (these cells are also known as i^3^Neurons) were thawed onto vitronectin coated 6 well tissue culture plates in E8 culture medium supplemented with 10 uM ROCK inhibitor. The next day, the E8 medium with ROCK inhibitor was removed and replaced with E8 medium without ROCK inhibitor. The cells were expanded and passaged using 500uM EDTA until colonies reached a confluency of 70-80% in 10cm tissue culture plates. Once at 70-80% confluence, iPSCs were split using Accutase to get a single cell suspension and 2-2.5×10^7^ cells were transferred onto Matrigel coated 15cm tissue culture dishes containing Neuronal Induction Media with doxycycline (see Supplementary table 4 for details). Cells were grown at 37°C and Neuronal Induction Media with doxycycline was changed daily for 3 days. After 3 days of induction, neurites were clearly visible, and the neurons were split using accutase and replated onto PDL coated XonaChip Imaging Slides (Fisher Scientific NC1648769) at a density of 1-5×10^5^ cells/mL in Neuronal Culture Media (see Supplementary Table 4 for details). Cells were then allowed to grow for 7 days prior to experimentation with Neuronal Culture Media changed every 3 days. During this period, cells were checked daily under a phase-contrast microscope to ensure neuronal health.

To stress cells, media was removed and replaced by Neuronal Culture Media containing the appropriate concentration of prostaglandin J2, E2, J2 + E2 (5uM or 10uM). After 15 hours, the cells were fixed with 4% PFA and IF performed as described above using the anti-SRRM2 antibody and DAPI. For the 0.6M Sorbitol condition cells were treated for 1 hour rather than 15 hours.

### siRNA transfections

For siRNA validation, HEK293 tau biosensor cells were plated at a density of 7.0×10^5^ cells per 6-well. The following day cells were transfected with 25 pmol siRNA with Lipofectamine RNAiMAX and collected 60 hours post-transfection. Cell pellets were lysed in 2X SDS loading buffer and processed for Western blotting as described above.

For flow cytometry HEK293 tau biosensor cells were transfected with siRNA as described. 24 hours post-transfection cells were plated at a density of 1.25×10^5^ cells/ 24-well. 24 hours later cells were seeded with tau brain homogenate for 24 hours prior to analysis. siRNAs used are detailed in Supplemental table 5.

### Flow cytometry

For siRNA experiments, HEK293T tau biosensor cells were transfected with siRNAs 48 hours prior to seeding with tau brain homogenate at a final concentration of 0.5ng/μl. For overexpression experiments, HEK293T tau biosensor cells were transfected with plasmids 24 hours prior to seeding with tau brain homogenate (0.5 ng/μl) and addition of TMRDirect Halo ligand (200nM) to label exogenously expressed Halo-tagged fusion proteins. 24-hours post seeding, cells were trypsinized, washed with PBS and filtered with 50um nylon mesh filters prior to cell sorting. Sorting was performed with a BD FACSCelesta^TM^ Cell Analyzer using the following filter sets: 561-585 (Halo), 405-450 (CFP) and 405-525 (FRET). Analysis was performed using FlowJo. Gating was performed in sequential steps, first sorting for cells, single cells, then (when applicable) gating based on Halo expression was performed. Lastly, gating for FRET+ cells was performed based on mock seeded cells to set a false FRET percentage at 1 as previously detailed (Furman et al., 2015). Integrated FRET Density was calculated as a product of the percentage of FRET positive cells and median fluorescence intensity.

### Image analysis using Ilastik and CellProfiler

To measure enrichment of endogenous SRRM2, C-terminal fragments, polyserine and controls in tau aggregates, images were first segmented into cytoplasmic tau aggregates, nuclear tau aggregates, nucleus, cytosol, and background using Ilastik with a minimum of five training images per condition per experiment. Images were hand annotated to show the location of the desired structures, which enabled the construction of a model that could then segment subsequent images. The original image and the image segmentation masks created by Ilastik were then used as inputs for a CellProfiler pipeline that calculated pixel intensity values of the 488 and 647 channels within the masked compartments. These compartmental measurements were used to calculate enrichment of SRRM2 in tau aggregates per image as follows: cytoplasmic tau aggregate enrichment = mean intensity within cytoplasmic tau aggregates per image / mean intensity within the cytosol per image.

To measure enrichment of endogenous PNN or its polyserine rich sequence and controls in tau aggregates, a CellProfiler pipeline was generated to segment the cytoplasm and nuclei of individual cells. Mean intensity measurements were taken and the fold enrichment reported for each cell as the mean intensity of Halo signal within tau aggregates / mean intensity of the remainder of the cytoplasm. This method was also used to quantify the fold enrichment of mRuby2 or mRuby2-G3BP1 signal in tau aggregates.

---

**Table S2.** Antibodies

| Antibody name | Manufacturer | Catalog # |
| --- | --- | --- |
| Rabbit anti-SRRM2 | ThermoFisher | PA5-66827 |
| Rabbit anti-PNN (Pinin) | ThermoFisher | 18266-1-AP |
| Rabbit anti-SETD1A | Millipore Sigma | HPA058376 |
| Rabbit anti-Halo | Promega | G9281 |
| Mouse anti-Tau-5 | Thermo Fisher | AHB0042 |
| Rabbit anti-B-III Tubulin | Cell Signaling | 5568T |
| Rabbit anti-MSUT2 | Sigma-Aldrich | HPA049798 |
| GAPDH-HRP | Santa Cruz | sc-47724 HRP |

**Table S3.**
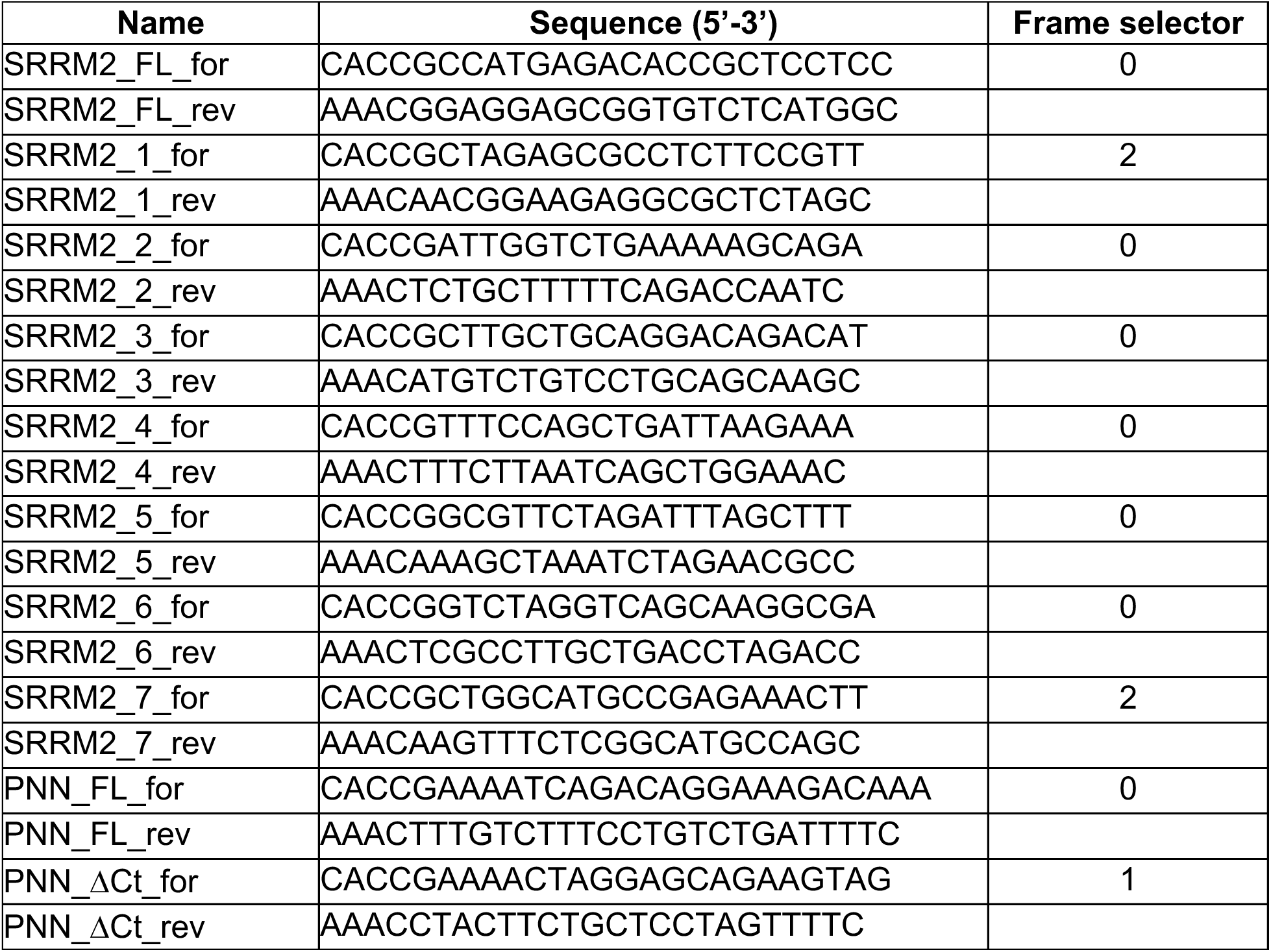
CRISPaint sgRNA Sequences

**Table S4.**
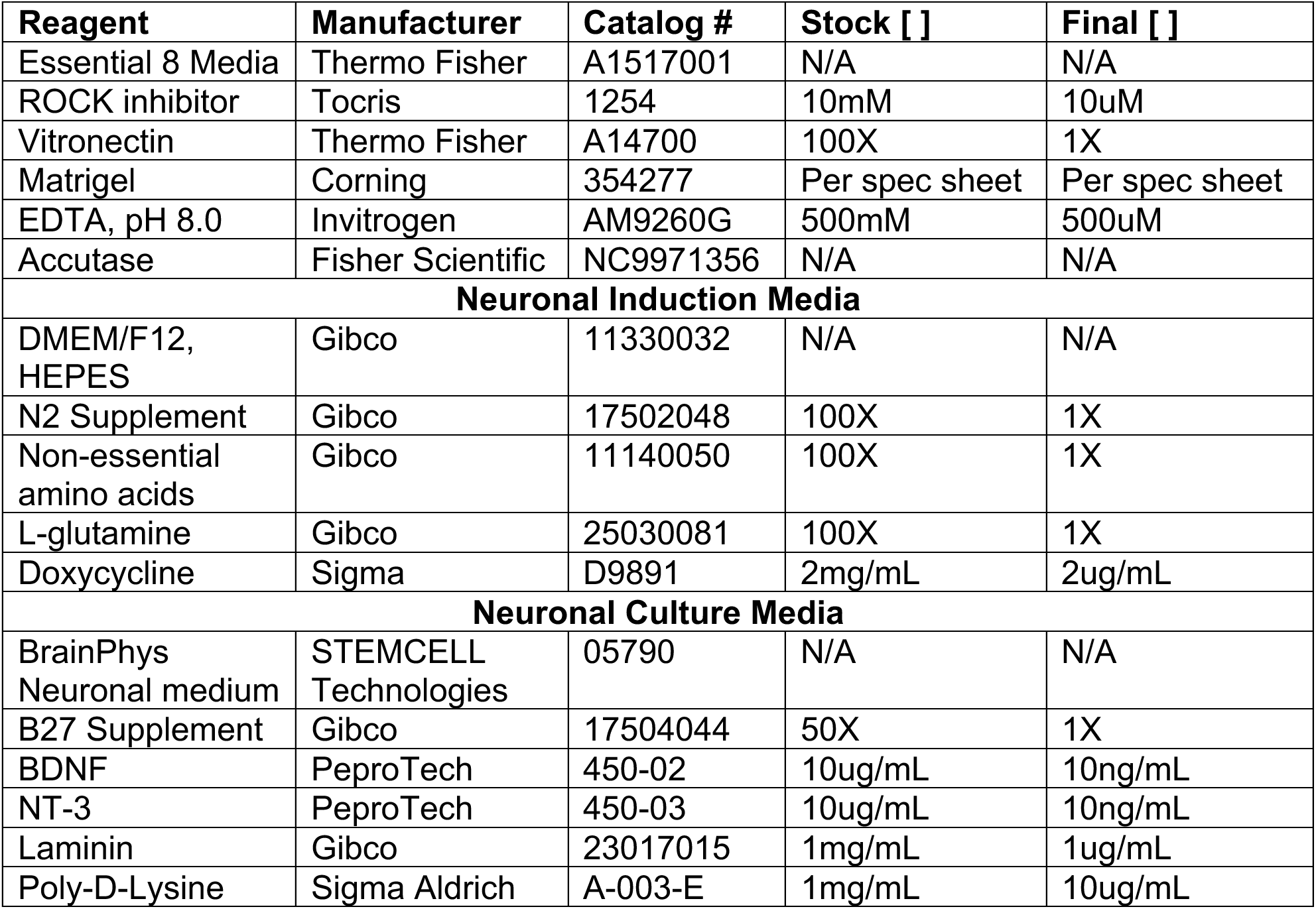
iPSC/Neuronal culture reagents

| Reagent | Manufacturer | Catalog # | Stock [ ] | Final [ ] |
| --- | --- | --- | --- | --- |
| Essential 8 Media | Thermo Fisher | A1517001 | N/A | N/A |
| ROCK inhibitor | Tocris | 1254 | 10mM | 10uM |
| Vitronectin | Thermo Fisher | A14700 | 100X | 1X |
| Matrigel | Corning | 354277 | Per spec sheet | Per spec sheet |
| EDTA, pH 8.0 | Invitrogen | AM9260G | 500mM | 500uM |
| Accutase | Fisher Scientific | NC9971356 | N/A | N/A |
| **Neuronal Induction Media** |  |  |  |  |
| DMEM/F12, HEPES | Gibco | 11330032 | N/A | N/A |
| N2 Supplement | Gibco | 17502048 | 100X | 1X |
| Non-essential amino acids | Gibco | 11140050 | 100X | 1X |
| L-glutamine | Gibco | 25030081 | 100X | 1X |
| Doxycycline | Sigma | D9891 | 2mg/mL | 2ug/mL |
| **Neuronal Culture Media** |  |  |  |  |
| BrainPhys Neuronal medium | STEMCELL Technologies | 05790 | N/A | N/A |
| B27 Supplement | Gibco | 17504044 | 50X | 1X |
| BDNF | PeproTech | 450-02 | 10ug/mL | 10ng/mL |
| NT-3 | PeproTech | 450-03 | 10ug/mL | 10ng/mL |
| Laminin | Gibco | 23017015 | 1mg/mL | 1ug/mL |
| Poly-D-Lysine | Sigma Aldrich | A-003-E | 1mg/mL | 10ug/mL |

**Table S5.** siRNA

| Target | Manufacturer | Catalog/siRNA ID # |
| --- | --- | --- |
| Control | Thermo Fisher | CAT# 4390843 |
| SRRM2 | Thermo Fisher | s24003 |
| MSUT2 | Millipore | EHU149521 |
| PNN (1) | Thermo Fisher | s10758 |
| PNN (2) | Thermo Fisher | s10759 |

**Figure S1:**
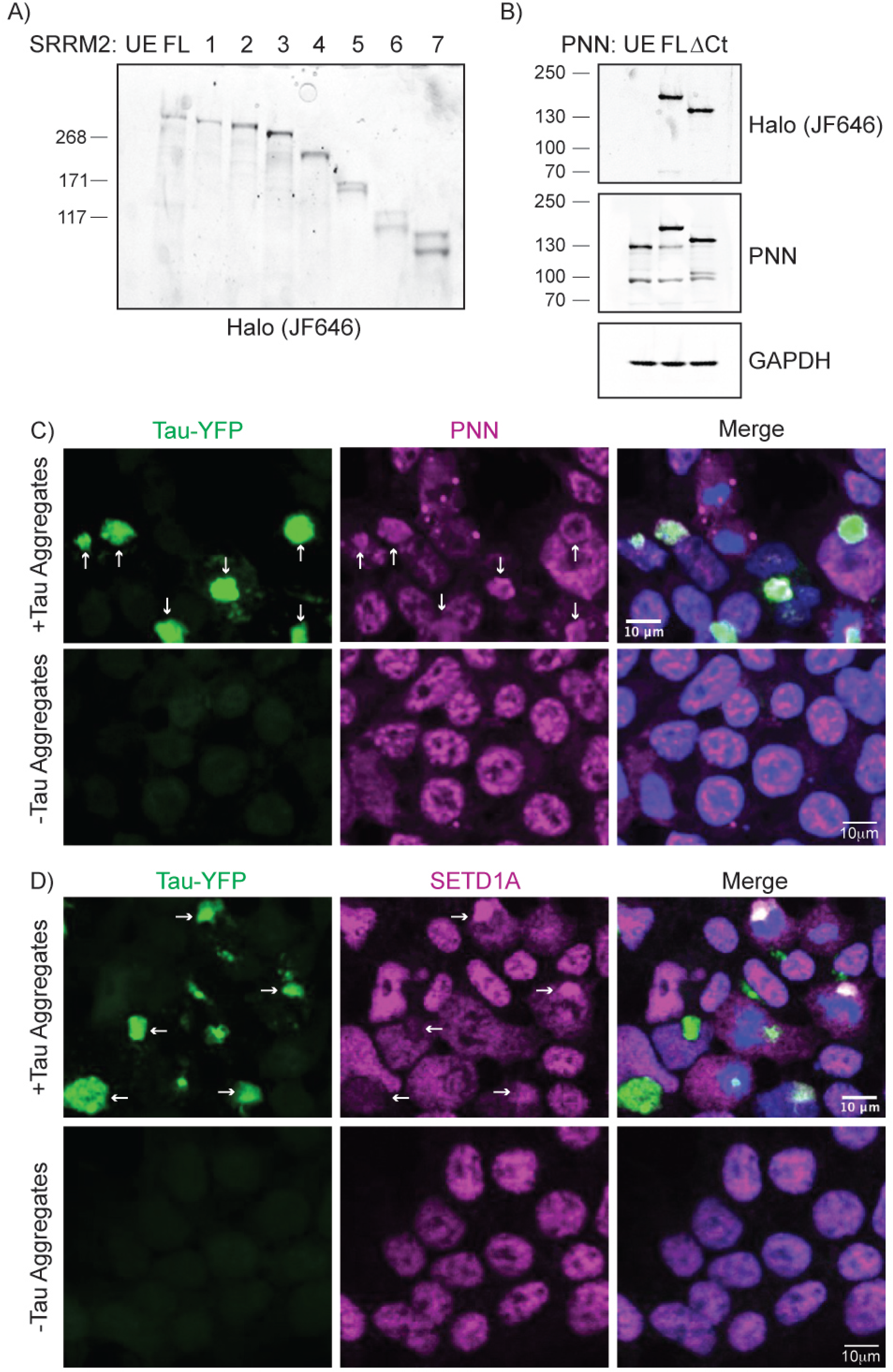
Polyserine-containing proteins enrich in tau aggregates. **(A)** Fluorescent imaging of Halo-tagged proteins from cell extracts of each SRRM2 truncation cell line. JF646 Halo ligand was added to cells prior to lysis and denaturation. **(B)** Fluorescent imaging of Halo-tagged proteins and Western blot for PNN and GAPDH from cell extracts of unedited, full-length (PNN_FL) and C-terminal truncated (PNN_ΔCt) cell lines. **(C)** Immunofluorescence of Tau-YFP (green), PNN (magenta), and DAPI (blue) in HEK293 biosensor cells with or without lipofection of clarified brain homogenate from tau transgenic mice (Tg2541). Tau aggregates colocalizing with PNN are denoted (white arrows). **(D)** Immunofluorescence of Tau-YFP (green), SETD1A (magenta), and DAPI (blue) in HEK293 biosensor cells with or without lipofection of clarified brain homogenate from tau transgenic mice (Tg2541). Tau aggregates colocalizing with PNN are denoted (white arrows).

**Figure S2:**
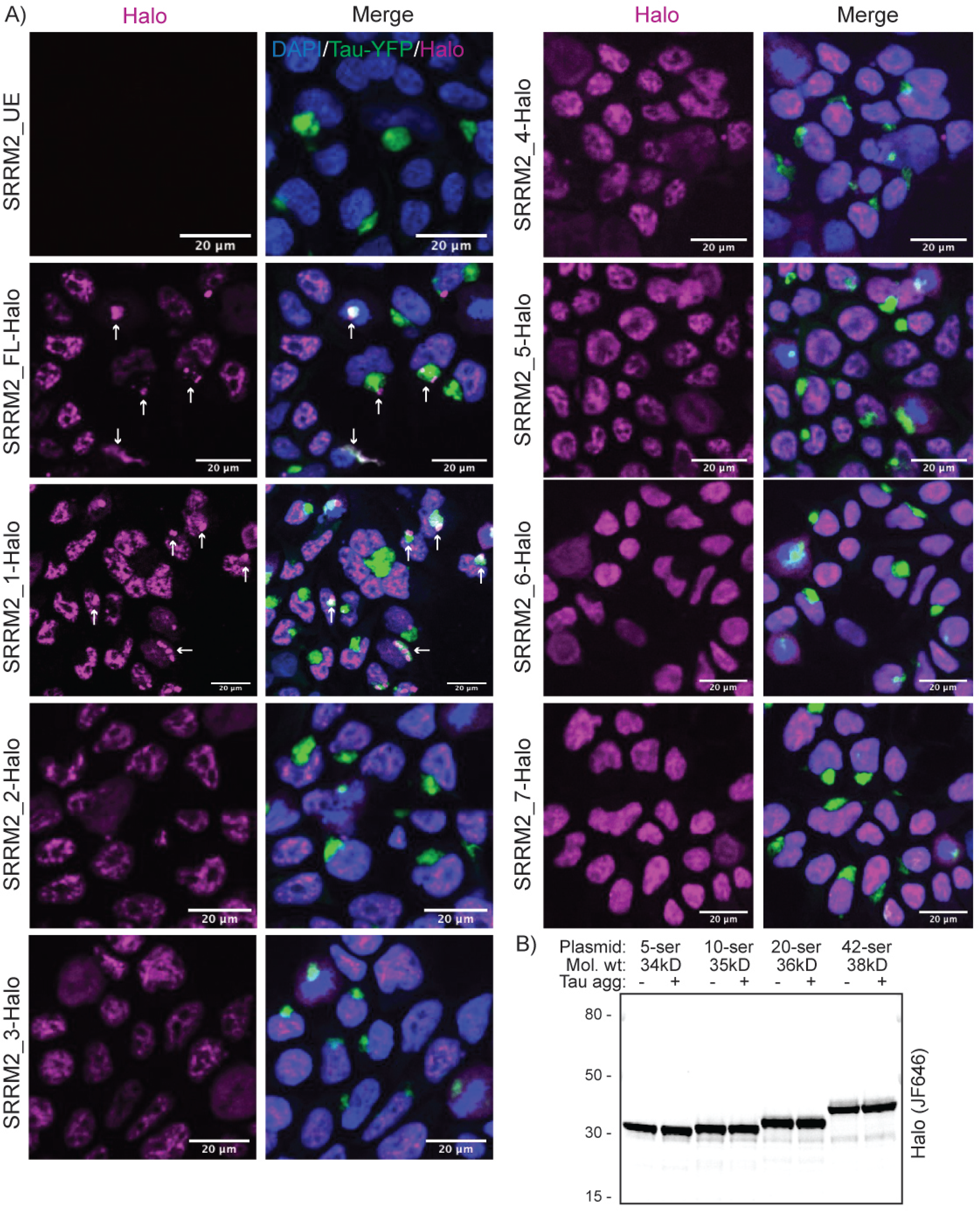
Characterization of Halo-tagged SRRM2 truncation cell lines. **(A)** Immunofluorescence of Halo (magenta), tau-YFP (green) and DAPI (blue) in Halo tagged SRRM2 truncation cell lines and SRRM2_UE (unedited, no Halo tag). White arrows show colocalization in SRRM2_FL-Halo and SRRM2_1-Halo. No colocalization observed in the other truncations. **(B)** Fluorescent imaging of an SDS-PAGE gel of Halo-tagged proteins in HEK293 cell lysate transfected with 5, 10, 20, or 42 polyserine-Halo constructs with or without transfection of clarified tau brain homogenate.

**Figure S3:**
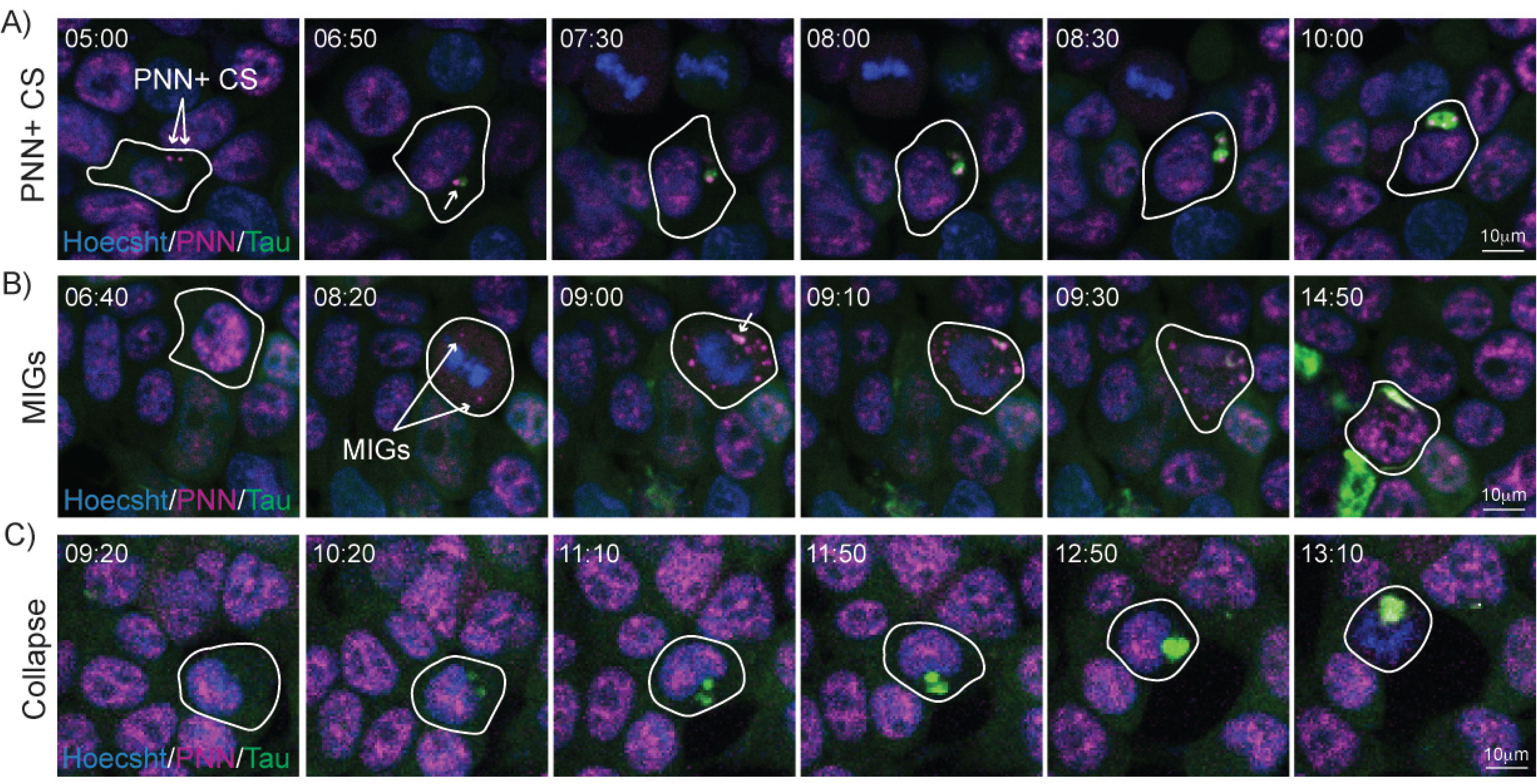
SRRM2 and PNN cytoplasmic granules are sites of tau aggregation. **(A-C)** Live imaging of Hoechst (blue), Tau-YFP (green) and PNN_FL-Halo (magenta) in HEK293 tau biosensor cells seeded with tau aggregates and monitored for 24 hours in 10-minute increments. Stills from live imaging display tau aggregate formation at PNN+ cytoplasmic speckles (A) (Movie 6), mitotic interchromatin granules (B) (Movie 7) and aggregate formation followed by nuclear collapse (C) (Movie 8). Time since the onset of imaging is displayed.

**Figure S4:**
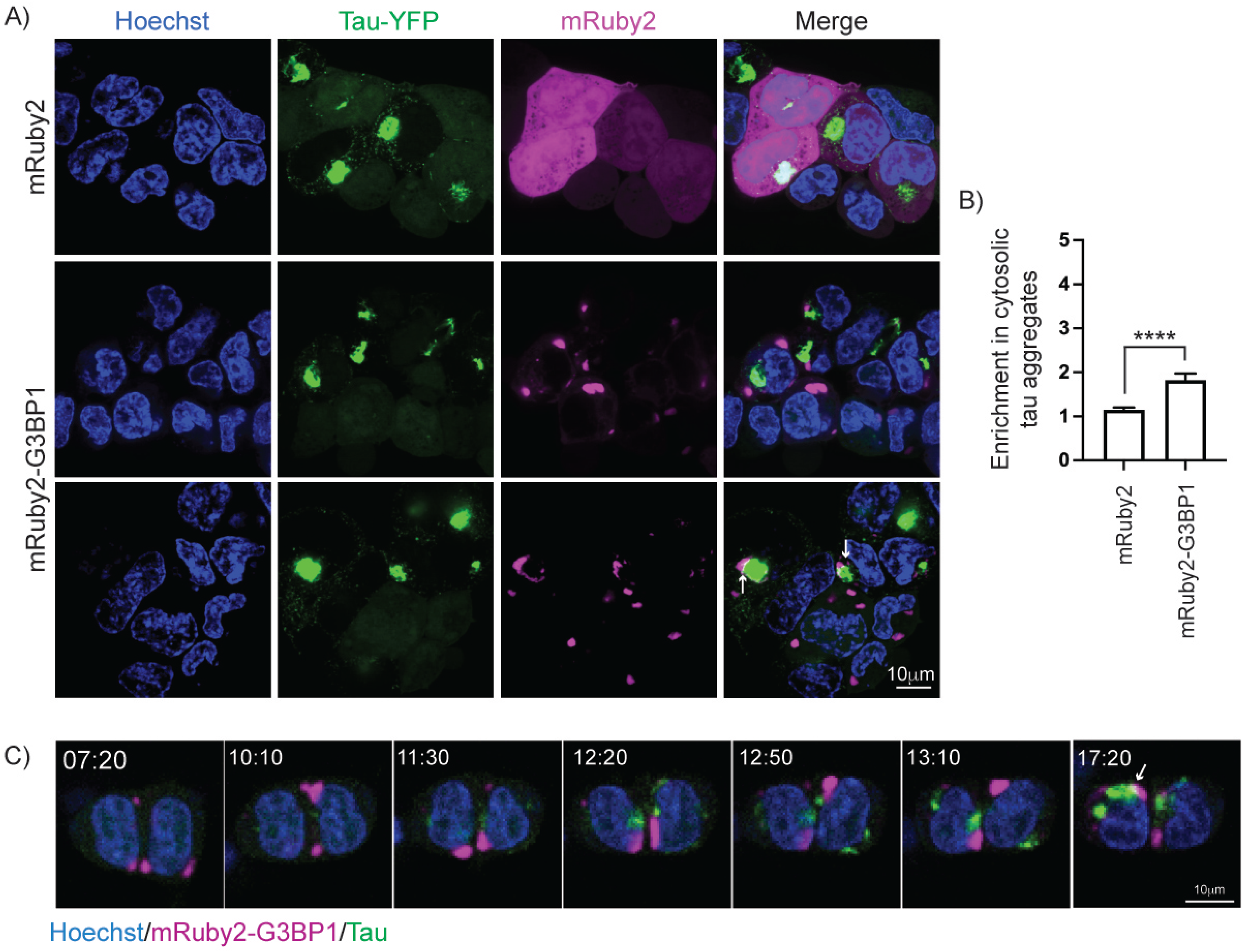
Stress granules are not preferred sites of tau aggregate formation. **(A)** Immunofluorescence of Hoechst (blue), Tau-YFP (green) and mRuby2 (magenta) in HEK293 tau biosensor cells stably transduced to express mRuby2 or mRuby2-G3BP1 30 hours post tau seeding and 24 hours after treatment with 50nM PatA. Docking of stress granules and tau aggregates are denoted by white arrows. **(B)** Quantification of the fold enrichment of mRuby2 in tau aggregates in HEK293 tau biosensor cells as in (A). Data represent mean and 95% CI. n>503 cells per group from three biological replicates. Statistics performed with Mann-Whitney test. (****) P < 0.0001. **(C)** Live imaging of Hoechst (blue), Tau-YFP (green) and mRuby2 (magenta) in HEK293 tau biosensor cells as in (A). Formation of stress granules and tau aggregates are independent with docking (white arrows) observed at later timepoints.

**Figure S5:**
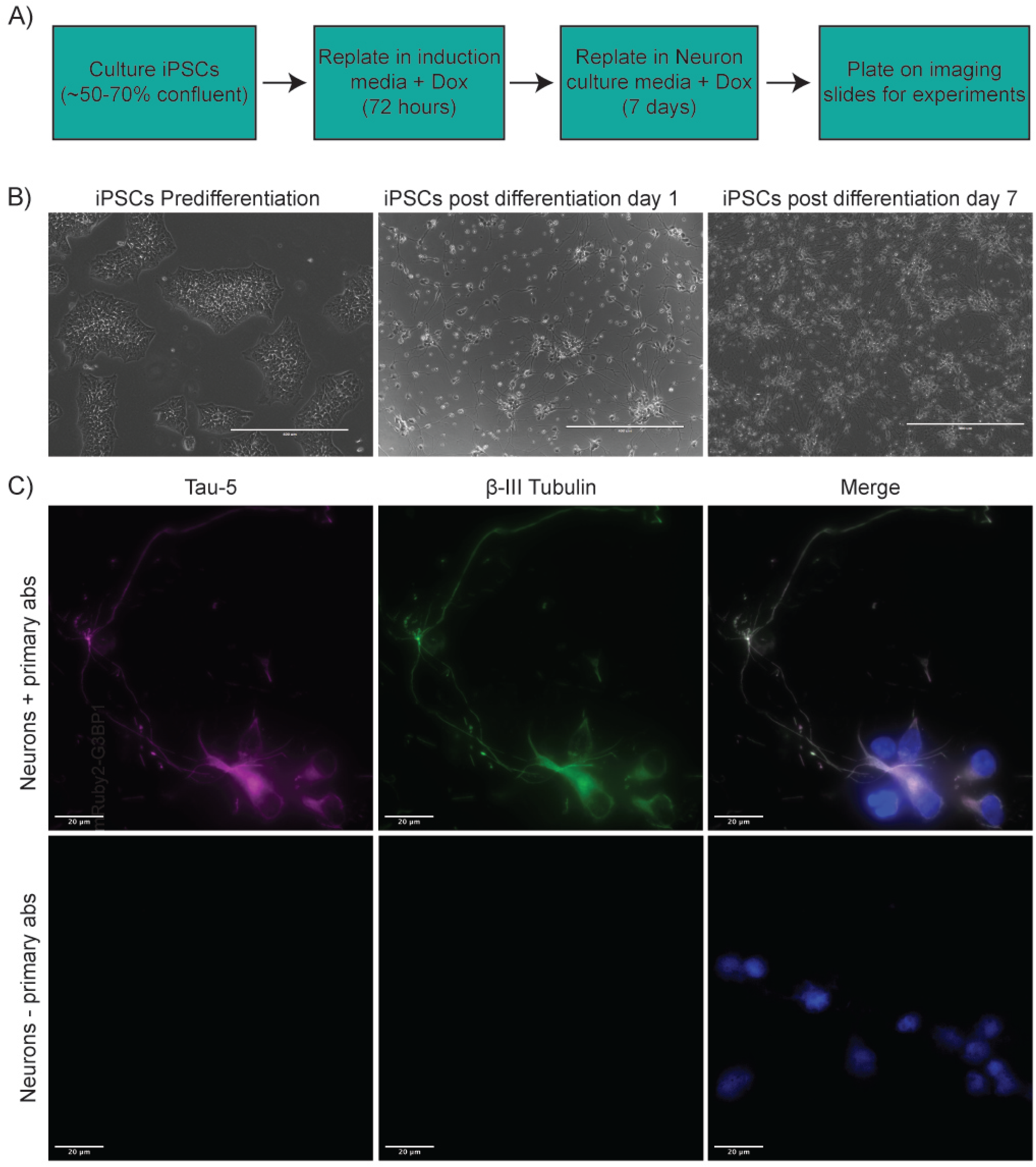
Characterization and validation of iPSC-derived neurons. **(A)** Differentiation protocol schematic for iPSC-derived iNeurons. **(B)** Brightfield images of iPSCs pre-differentiation, at day 1 post-differentiation, and day 7 post-differentiation. **(C)** Immunofluorescence of neuronal markers β-III Tubulin (green) and Tau-5 (magenta) and DAPI (blue) in iNeurons at 8 days post differentiation.

**Figure S6:**
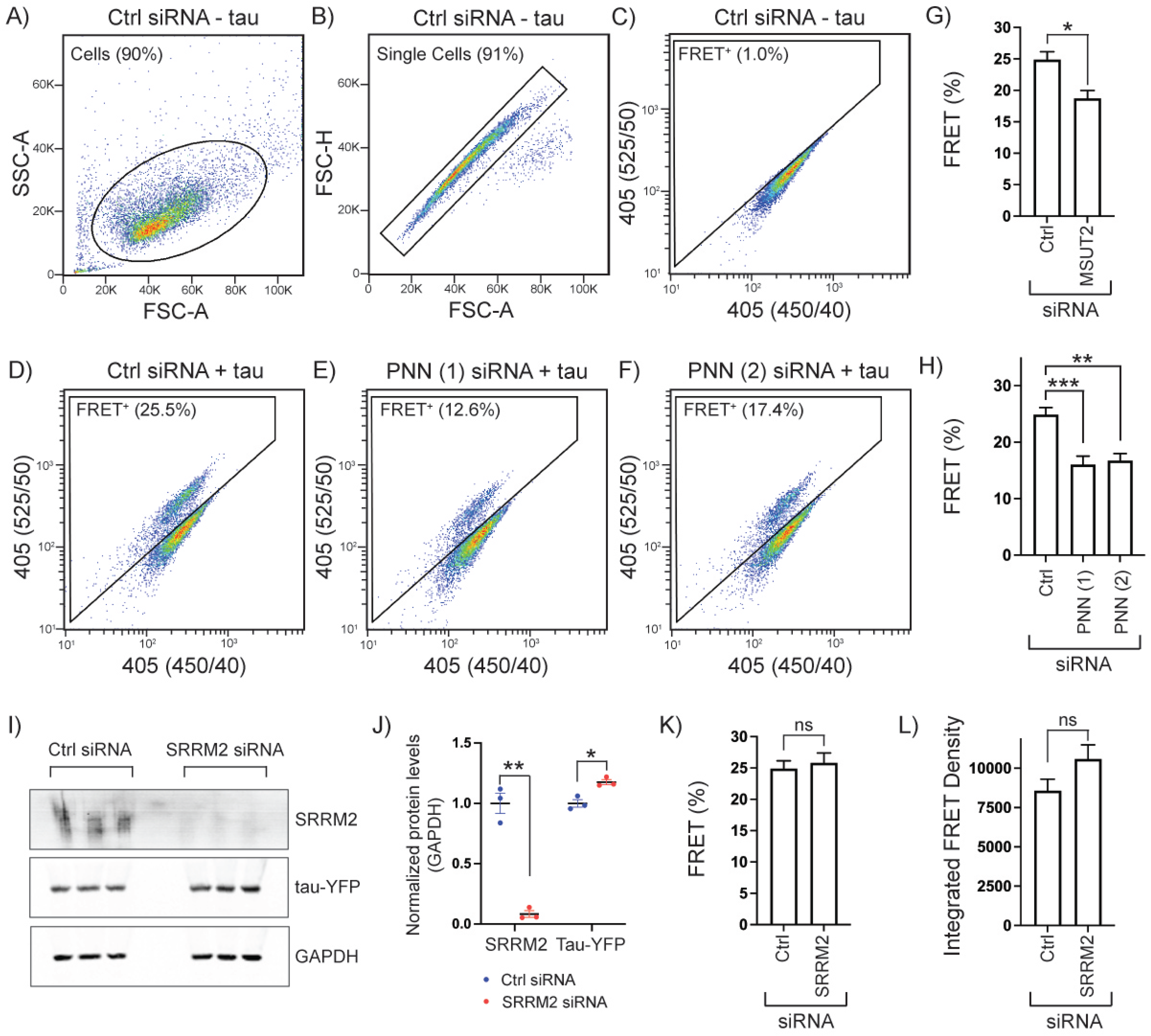
MSUT2 and PNN knockdown reduces tau aggregation by FRET-based flow cytometry. **(A)** Plot of forward scatter area (FSC-A) versus side scatter area (SSC-A) with representative gating for HEK293 biosensor cells treated with control siRNA. **(B)** Scatter plot of FSC-A versus forward scatter height (FSC-H) for cells selected in (A) with representative gating for single cells of HEK293 biosensor cells treated with control siRNA. (**C)** Scatter plot of CFP [405 (450/40)] versus FRET [405 (525/50)] signal in HEK293 biosensor cells treated with control siRNA selected in (B) with representative gating for FRET+ cells. **(D-F)** Scatter plot of CFP [405 (450/40)] versus FRET [405 (525/50)] signal in HEK293 biosensor cells treated with control or PNN siRNAs and seeded with clarified tau brain homogenate from populations as selected in (B) with representative gating for FRET+ cells. **(G)** Percentage of FRET+ HEK293 biosensor cells treated with control or MSUT2 siRNA measured by flow cytometry and analyzed as in (Figure S6A-C). Data represent mean and SEM. Statistics performed with Mann-Whitney test. (*) P < 0.05. **(H)** Percentage of FRET+ HEK293 biosensor cells treated with control of each PNN siRNA measured by flow cytometry and analyzed as in (A-F). Data represent mean and SEM. Statistics performed with one-way ANOVA. (**) P < 0.01; (***) P < 0.001. **(I)** Western blot of SRRM2, tau-YFP and GAPDH protein levels in HEK293 biosensor cells treated with control or SRRM2 siRNA. **(J)** Quantification of Western blot shown in (I) normalized to GAPDH. Bars represent mean and SEM. Statistics performed with unpaired t-test with Welch’s correction. **(K)** Percentage of FRET+ HEK293 biosensor cells treated with control or SRRM2 siRNA measured by flow cytometry and analyzed as in (Figure S6A-C). Data represent mean and SEM. Statistics performed with Mann-Whitney test. (ns) P > 0.05. **(L)** Integrated FRET Density of HEK293 biosensor cells treated with control or SRRM2 siRNA measured by flow cytometry and analyzed as in (Figure S6A-C). Data represent mean and SEM. Statistics performed with Mann-Whitney test. (ns) P > 0.05.

**Figure S7:**
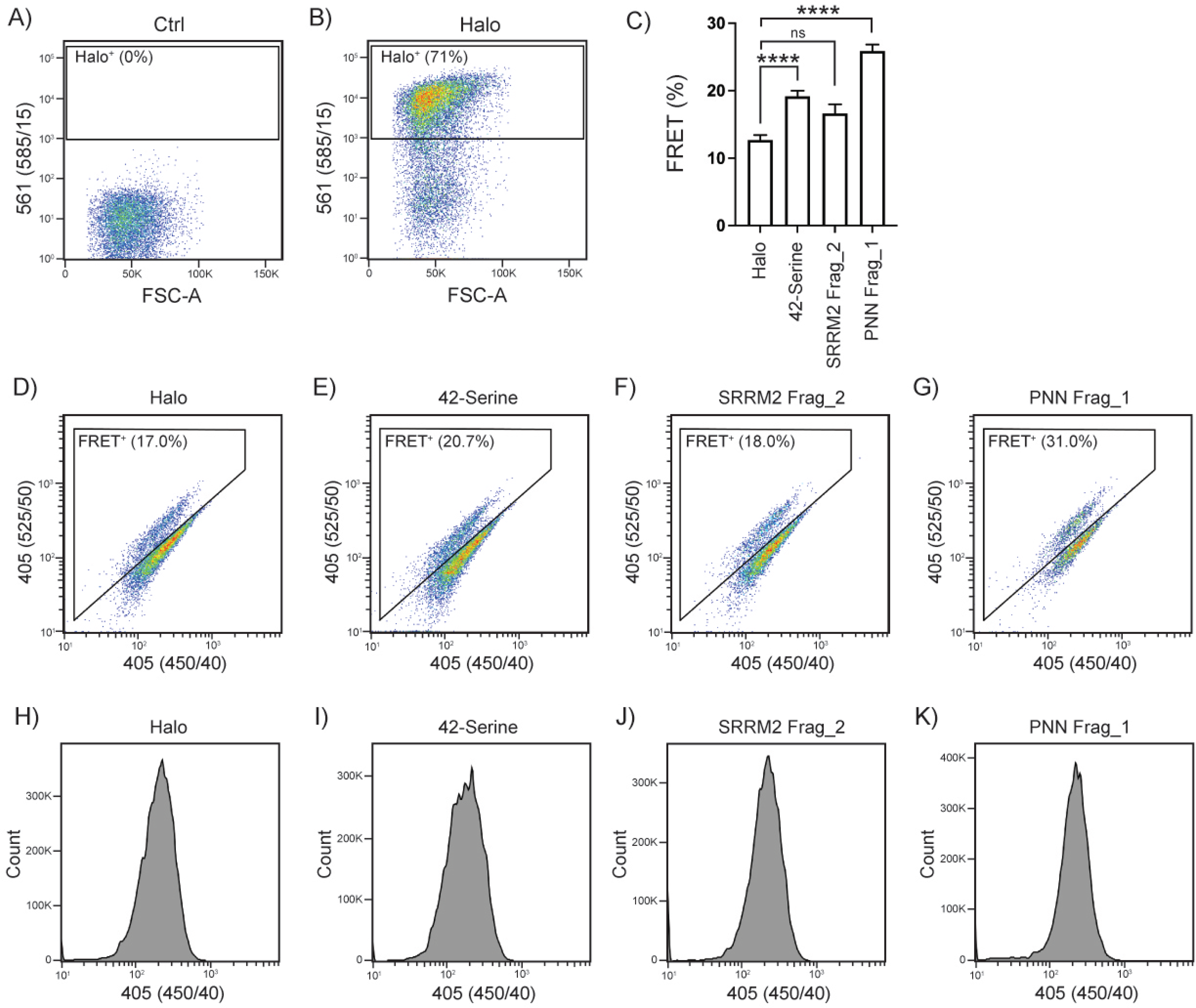
Overexpression of polyserine does not alter tau levels. **(A, B)** Scatter plot of FSC-A versus Halo intensity signal [561 (585/15)] in HEK293 biosensor cells mock transfected or transfected with Halo showing representative gating for Halo+ cells performed following gating steps detailed in Figure S6A,B. **(C)** Percentage of FRET+ HEK293 biosensor cells transfected with Halo, 42-Serine, SRRM2 Frag_2 and PNN Frag_1 constructs by flow cytometry and analyzed by gating in Figure S6A,B, subsequently Figure S7A,B and lastly for FRET positivity as in Figure S6C. Data represent mean and SEM. Statistics performed with one-way ANOVA. (ns) P > 0.05; (****) P < 0.0001. **(D-G)** Scatter plots of CFP [405 (450/40)] versus FRET [405 (525/50)] signal in HEK293 biosensor cells transfected with Halo, 42-Serine, SRRM2 Frag_2 and PNN Frag_1 after gating for cells and single cells as demonstrated in Figure 6A-B with representative gating for FRET+ cells. **(H-K)** Histograms of CFP [405 (450/40)] signal in HEK293 biosensor cells transfected with Halo, 42-Serine, SRRM2 Frag_2 and PNN Frag_1 after gating for cells and single cells.

